# Prefrontal coding of learned and inferred knowledge during REM & NREM sleep

**DOI:** 10.1101/2021.04.08.439095

**Authors:** Kareem Abdou, Mohamed H. Aly, Ahmed Z. Ibrahim, Kiriko Choko, Masanori Nomoto, Reiko Okubo-Suzuki, Shin-ichi Muramatsu, Kaoru Inokuchi

## Abstract

Idling brain activity has been proposed to facilitate inference, insight, and innovative problem-solving^1-4^. However, it remains unclear how and when the subconscious brain can create novel ideas. Here, we show coordinated roles of non-rapid eye movement (NREM) and rapid-eye movement (REM) sleep in weaving inferential information from learned information. In a transitive inference paradigm, mice gained the inference one day, but not shortly, after complete training. Inhibiting the neuronal computations in the anterior cingulate cortex (ACC) during post-learning either NREM or REM sleep, but not wakefulness, disrupted the inference without affecting the learned knowledge. *In vivo* Ca^2+^ imaging showed gradual development of inference-related representation, peaking at post-training REM sleep. Neurons representing learned information co-reactivated significantly during post-training NREM sleep, while their co-reactivation with neurons representing inferential information was significantly higher during REM sleep. Furthermore, after insufficient learning, artificial activation of medial entorhinal cortex-ACC dialogue during only REM sleep created inferential knowledge. These findings suggest that NREM sleep organizes the scattered learned knowledge in a complete hierarchy, while REM sleep computes the inferential information from the organized hierarchy. Collectively, our study provides a mechanistic insight on NREM and REM coordination in weaving inferential knowledge, thus highlighting the power of idling brain in cognitive flexibility.

## Introduction

Cognitive flexibility is a distinctive feature that is required for higher learning^5^. Inferential reasoning is a prominent property of cognitive flexibility since it relies on flexible and systematic organization of existing knowledge^3,5^. Activation patterns of neurons in the medial prefrontal cortex (mPFC) have been suggested to represent inferential reasoning^6,7^.

Experience-related neural representations are re-expressed during post-learning awake rest and sleep periods^8,9^. This neural replay has been proposed to consolidate memories^8,10-13^. Previous work has found that awake replay events that occur during a spatial task not only represent recent experiences, but also novel spatial paths that might be confronted, which indicates that a whole spatial map has been formed^14-16^. Targeted reactivation of an implicitly learned sequence memory during sleep enables the development of explicit awareness of that sequence^17^. Previous reports have shown an offline improvement in a word-pair association task following sleep^18-21^.

Enhanced cognitive performance has been linked with offline activity during non-rapid eye movement (NREM)^22^ and rapid eye movement (REM)^23^ sleep, but the most pertinent sleep stage and the circuits involved are still unclear. REM and NREM sleep have unique neurophysiological processes that impact cognition^24,25^. REM sleep is characterized by ponto-geniculo-occipital waves, theta rhythm, high acetylcholine levels, and upregulation of plasticity-related genes^26^. Long-term potentiation is easily induced during REM sleep^27^. NREM sleep is characterized by regular occurrence of slow wave delta activity, high protein synthesis levels, and a rise in depotentiation-related genes and drop in long-term potentiation-related genes^24,28-33^. Therefore, sleep stages differentially affect neuronal functions and may therefore modify cognitive capabilities. Furthermore, previous reports have proposed that NREM and REM sleep have differential contribution in several brain functions; for example, memory consolidation^34^, visual learning^35^, skills and schemas^36^. Consequently, we hypothesize that NREM and REM sleep have distinct roles in the processing of learned and inferential information.

The aforementioned studies^14-17,22,23^ highlight the involvement of online and offline brain activity in cognitive flexibility. However, the temporal, cellular and circuit mechanisms underlying the emergence of inference from chronologically separated, yet overlapping, experiences are still unclear.

## Results

### Randomizing existing knowledge is necessary for the emergence of inferential behaviour

We developed a transitive inference paradigm in mice that assesses the ability to infer new information that was not learned before, based on previous knowledge of overlapping memories. First, food-restricted mice were allowed to explore and habituate to an arena consisting of five different contexts (Context A, B, C, D, and E) (Fig. 1a). These contexts were different in terms of geometry and/or floor texture and/or wall color and pattern. During the habituation period, mice did not show a significant innate preference to any context (Extended Data Fig. 1). Then, they were trained on a series of two-context discrimination trials called premise pairs (A > B, B > C, C > D, and D > E; where “>” means that the former context is more preferred than the latter one). Mice should prefer the context in which they were rewarded with a sucrose tablet. Every day, mice were trained on two premise pairs in two sessions, with each session consisting of five trials. During each trial, mice were put into the arena with only two opened contexts, while the other three contexts were closed (Fig. 1b). After staying in the assigned context for a consecutive 10 sec, mice received a sucrose tablet (Extended Data Video 1). After reaching the criterion for correct performance (80% correct trials) in the four premise pairs (Fig. 1c), mice were divided into two groups. The first group was subjected to reinforcement sessions only, while the second group received randomized training sessions in which each pair was presented with a different pair every day (Fig. 1b and Methods). Then, mice were subjected to inference (B & D) and non-inference (A & E) tests on the last day of training (T1) and in the subsequent 3 days (T2, T3, and T4). Contexts B and D had an equal valence during training as they were the preferred context in one premise pair (B > C and D > E, respectively), and were the avoided context in the other premise pair (A > B and C > D, respectively), such that the performance in the inference test would be purely due to inference from the hierarchy. On the other hand, context A was usually the preferred context, while context E was usually un-preferred; therefore, the performance during tests with the A and E contexts would not be due to the inference. Mice that received randomized training inferred correctly in all tests except for in T1, as shown by an increased number of correct trials and a decreased latency time to choose the correct context (Fig. 1d, e). On the other hand, mice that received reinforcement training only without randomization did not exhibit inference in any tests (Fig. 1d, e); however, they had successfully learnt the original premise pairs (Fig. 1c), which indicates that knowledge of the premise pairs did not assure the emergence of inference. This showed the necessity of randomization during acquisition of the premise pairs knowledge for gaining the inference. In both groups, the percent of correct trials was significantly higher than chance level (50%) in the non-inference test, and their memories of the original premises were intact (Extended Data Fig. 2a).

**Fig. 1.**
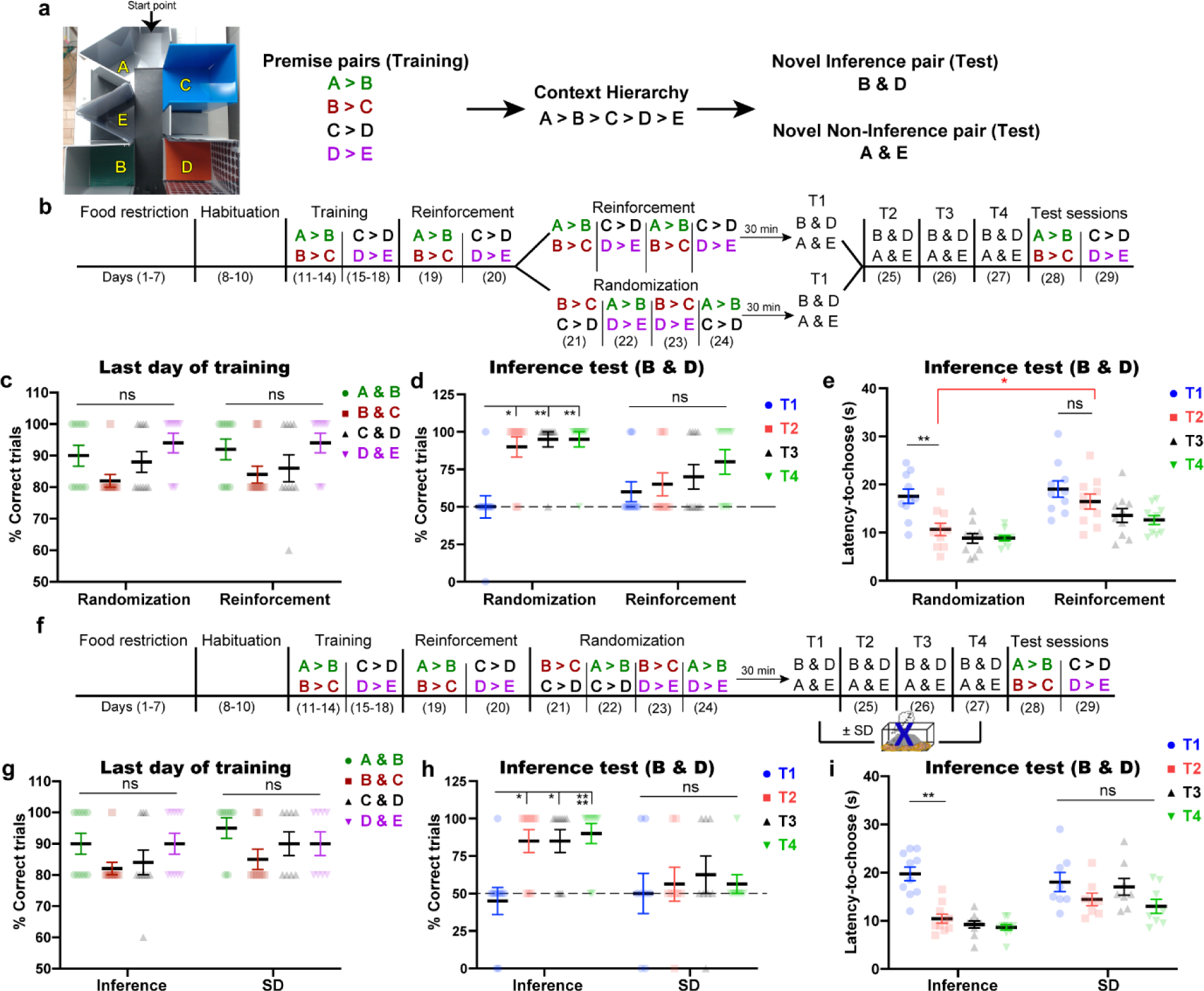
The emergence of inference requires sleep after training. **a,** Representative photo of the arena and the five different contexts with an arrow representing the starting point (left). The four premise pairs with the contexts’ hierarchy (right). **b,** The behavioural schedule used to establish the transitive inference paradigm in mice. The contexts’ hierarchy was as follows: A > B > C > D > E. The order of presentation of the premise pairs during the randomization stage is different across animals within the same group (See Methods; Table 1). **c,** Performance during the last day of training (day 14 for A > B and B > C; day 18 for C > D and D > E) for each premise pair was calculated as the percent of correct trials out of the total number of trials. **d-e,** Performance during test sessions; the percent of correct trials in the inference test (**d**), the latency time to choose in the inference test (**e**). **f,** Behavioural protocol for the transitive inference paradigm; SD, sleep deprivation. **g,** Percent of correct trials during the last day of training (day 14 for A > B and B > C; day 18 for C > D and D > E) for each premise pair. **h-i,** Performance during test sessions; percent of correct trials during the inference test (**h**), latency time to choose during the inference test (**i**). T1, test 1; T2, test 2; T3, test 3; T4, test 4. *n* = 10 mice/group; *n* = 8 mice for the sleep deprivation (SD) group. Statistical comparisons were made using a two-way repeated-measures ANOVA with Tukey’s multiple comparison test (**c-e, g-h**) and Holm-Sidak’s test between groups (**e**). \**P* < 0.05; \*\**P* < 0.01; \*\*\*\**P* < 0.0001; ns, not significant (*P* > 0.05). Data are presented as the mean ± standard error of the mean (s.e.m.).

### The emergence of inference requires sleep after training

The inability to infer correctly in T1 and the expression of inference in T2 suggest that inference does not arise online during the repeated training, but instead may require an incubation period or sleep following training. To examine which factor is critical for the inference process, we kept the incubation period intact (the same as before), but mice were sleep-deprived for 4 h directly after test sessions (Fig. 1f). Mice met the criterion for correct training performance, with a non-significant difference in the percent of correct trials across all premise pairs (Fig. 1g). Sleep deprivation blocked the inference; the correct performance of mice remained at chance level and their decision making was delayed (Fig. 1h, i). However, sleep deprivation did not affect performance on the non-inference test or the original premises’ tests (Extended Data Fig. 2b). This suggests that sleep after the last training is crucial for constructing the whole hierarchy of preferences and the subsequent development of inference, but that it is not essential for the A and E pair or for the maintenance of the original premises’ representations. The inferential behaviour that appeared in T2 in the inference group was not due to learning during T1 since mice were not provided with sucrose in T1, even when the correct choice was made.

To test whether the inference observed in T2 was facilitated by T1 exposure in which B and D pair was presented to mice, mice were exposed to the same behaviour protocol without exposure to T1 (Extended Data Fig. 3a). After successful training phase (Extended Data Fig. 3b), mice inferred correctly during subsequent test sessions (Extended Data Fig. 3c, d). This result indicates that T1 exposure is not necessary for the emergence of inference during T2, which supports that inference is a higher-order process than a result of direct learning.

### Organizing existing knowledge into a hierarchical order is necessary for inference

To confirm that mice gained the inference after arranging the premise pairs into a hierarchical order, we trained a group of mice in the same arena, with the same number of trials, but with premise pairs that did not form a complete hierarchy (A > B, B > C, E > D, D > C; Extended Data Fig. 4a, b). Mice learnt the premise pairs well and reached the criterion of correct performance (Extended Data Fig. 4c). Lacking the complete hierarchy led to failed inference (Extended Data Fig. 4d, e), although mice memorized the original premise pairs well (Extended Data Fig. 4g). Since context A and context E were always preferred in the premise pairs training, mice did not prefer either of these during the non-inference test (Extended Data Fig. 4f). These results demonstrate that extracting inference requires the organization of previously acquired knowledge into a hierarchical order.

### Prelimbic cortex activity is not required for the emergence of inference

The mPFC has been proposed to process the mental schema that is utilized to build an inference, and may also compute the inferred outcomes^6,7,37-39^. However, these proposals were derived from lesion and functional brain imaging studies, and lack the causality for temporal and topographic specificity that are necessary for inference development. To directly unravel these ambiguities, we manipulated the activity of mPFC sub-region, the prelimbic cortex (PL) during sleep and awake states. The PL of mice was injected with adeno-associated virus 9 (AAV9) encoding a light-sensitive neuronal silencer (ArchT 3.0) with enhanced yellow fluorescent protein under the control of calcium/calmodulin-dependent protein kinase II (CaMKII) to label excitatory neurons with ArchT 3.0 (Extended Data Fig. 5a). Then, mice were subjected to the transitive inference task, where optogenetic inhibition was performed during either wakefulness or sleep on the last training day and testing days (Extended Data Fig. 5a, b). Mice achieved the standard for correct performance in all premise pairs (Extended Data Fig. 5c, d). Inhibiting PL activity during sleep or wakefulness after the training did not affect the inference, non-inference test, or premise pairs memories (Extended Data Fig. 5e-h), which indicates that PL activities during sleep and awake states are not required for the formation of inference.

### Anterior cingulate cortex computations during sleep are crucial for the emergence of inference

Next, we manipulated another mPFC sub-region, the anterior cingulate cortex (ACC). Mice received injections into the ACC of AAV-CaMKII-ArchT 3.0-eYFP to label excitatory neurons with ArchT 3.0 (Fig. 2a). Afterwards, mice completed the transitive inference task, where optogenetic inhibition was performed during either wakefulness, NREM sleep, or REM sleep on the last training day and testing days (Fig. 2b, c). Mice learnt the premise pairs well, which was reflected by the high number of correct trials that reached the criterion for correct performance (Fig. 2d). Optogenetic inhibition of the ACC (Extended Data Fig. 6a) during wakefulness did not impact inference ability, as mice achieved a high correct performance during inference, non-inference, and premise pairs tests (Fig. 2e, f and Extended Data Fig. 2c). Conversely, mice that received optogenetic inhibition of the ACC either during NREM or REM sleep failed to infer correctly, and the percent of correct trials was not significantly different from chance level (Fig. 2e). Moreover, the latency to choose was significantly longer in the NREM and REM groups than that of the awake group (Fig. 2f). These data indicate that ACC computations are necessary for the evolution of inference ability during sleep, but not during wakefulness. However, this perturbation to ACC dynamics had no influence on the non-inference pairs or the maintenance of premises knowledge (Extended Data Fig. 2c).

**Fig. 2.**
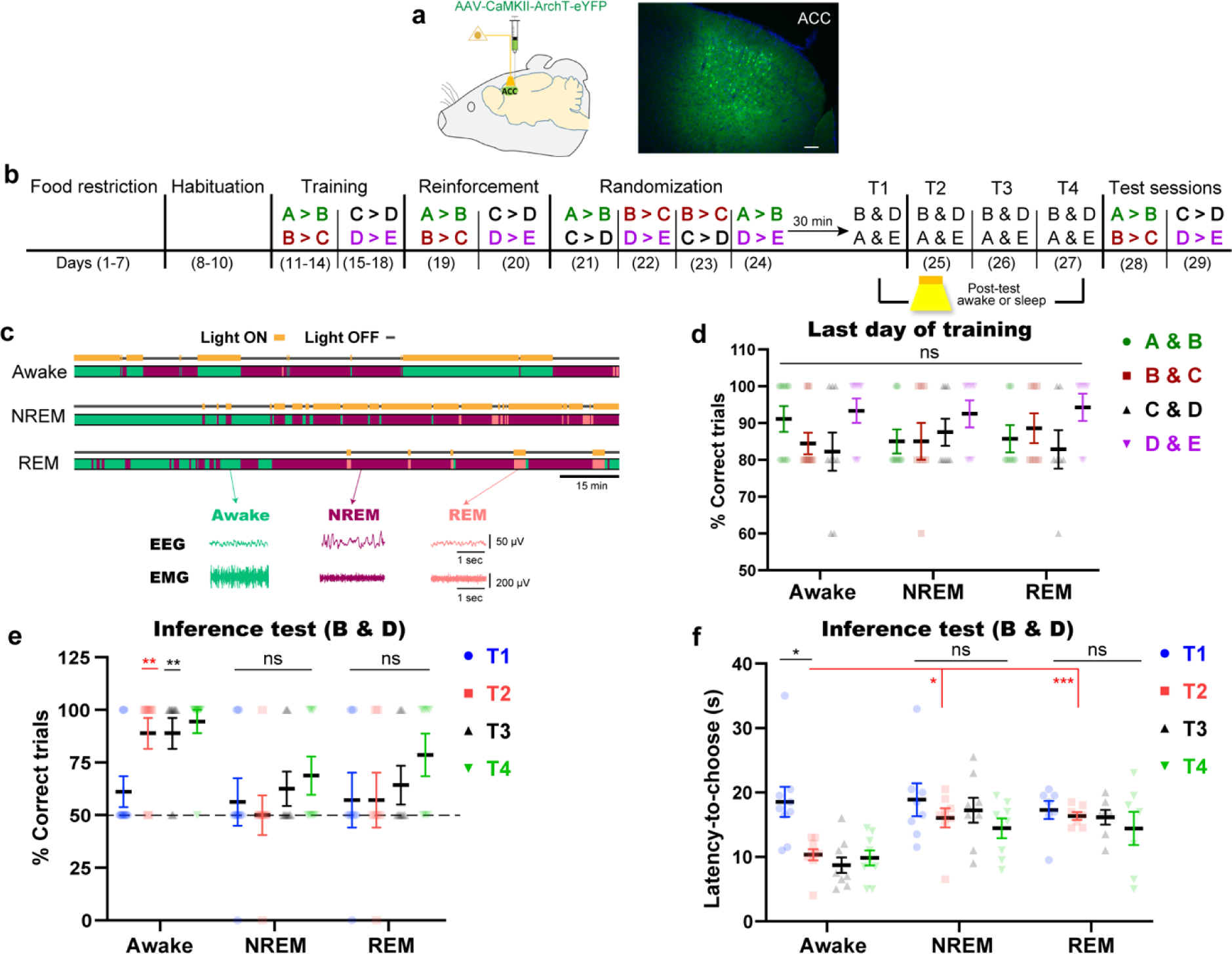
ACC computations during sleep are crucial for the emergence of inference. **a,** Labelling excitatory neurons of the ACC with ArchT (left), and the expression of ArchT-eYFP (green) in the ACC (right). Blue, 4′,6-diamidino-2-phenylindole (DAPI) staining. Scale bar, 100 µm. **b**, Behavioural schedule used to manipulate ACC activity during sleep and awake periods after test sessions. The order of presentation of the premise pairs during the randomization stage is different across animals within the same group (See Methods; Table 1). **c**, Diagram showing the state-specific manipulation (top) and representative electroencephalogram (EEG) and electromyography (EMG) (bottom) traces. Scale bar, 15 minutes. **d,** Performance during the last day of training (day 14 for A > B and B > C; day 18 for C > D and D > E) for each premise pair. **e-f,** Performance during test sessions; percent of correct trials during the inference test (**e**), latency time to choose during the inference test (**f**). T1, test 1; T2, test 2; T3, test 3; T4, test 4. *n* = 9 mice for the awake group; *n* = 8 mice for the non-rapid eye movement (NREM) sleep group; *n* = 7 mice for the rapid eye movement (REM) sleep group. Statistical comparisons were made using two-way repeated-measures analysis of variance (ANOVA) with Tukey’s multiple comparison test (**d-f**). In (**e**), the statistical significance denotes the comparison between performance relative to the chance level (50%). \**P* < 0.05; \*\**P* < 0.01; \*\*\**P* < 0.001; ns, not significant (*P* > 0.05). Data are presented as the mean ± standard error of the mean (s.e.m.).

### Inference representations develop gradually, peaking during REM sleep after training

To examine how the offline brain organizes the previously acquired knowledge and builds novel inferential information, we tracked the ACC neuronal activity via Ca2+ imaging. Wildtype mice were injected with AAV9-Syn-janelia-GCaMP7 into the ACC (Fig. 3a) and a custom-built electroencephalogram/electromyography (EEG/EMG) 5-pin system was installed into the skull, as previously described^40^, to record the neuronal dynamics during awake and sleep states (Fig. 3b). The same ACC neurons were tracked across sessions^8^ along the transitive inference task (Fig. 3c-e), using an automated sorting system^41^ to extract each neuron’s calcium (Ca2+) activity which were normalized to z-scores (see methods for details). Offline stages either awake or sleep were differentiated according to the automatically enumerated EMG root mean square value (ERMS), EEG delta power (1–4 Hz) RMS (dRMS), and EEG theta power (6–9 Hz) RMS, as described previously^40,42^. Sleep stages discrimination was concluded based on the delta/theta ratio value (Fig. 3b; see Methods for details).

**Fig. 3.**
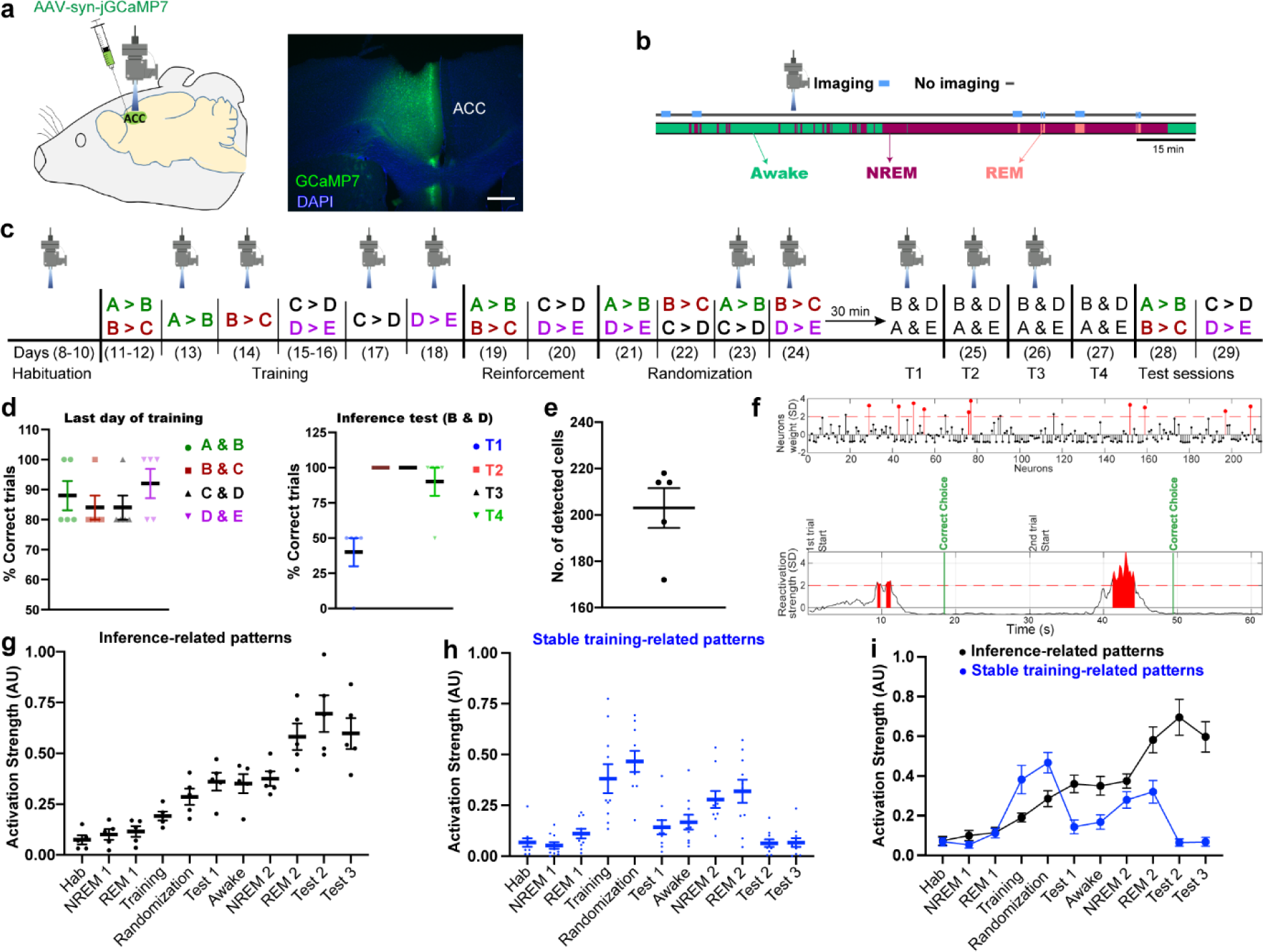
Identifying inference-related ensembles that emerge gradually and become evident during REM sleep. **a,** Labelling of the ACC with GCaMP7 (left), and the expression of GCaMP7 (green) in the ACC (right). Blue, 4′,6-diamidino-2-phenylindole (DAPI) staining. Scale bar, 100 µm. **b**, Diagram showing the state-specific imaging. **c,** The behavioural schedule used to capture the Ca^2+^ transients across the task. The order of presentation of the premise pairs during the randomization stage is different across animals within the same group (See Methods; Table 1). **d,** Performance during the last day of training (day 13 for A > B; day 14 for B > C; day 17 for C > D; day 18 for D > E) for each premise pair (left). Performance during test sessions (right). T1, test 1; T2, test 2; T3, test 3; T4, test 4. *n* = 5 mice. **e,** Total number of detected cells in each mouse. **f,** Example of ACC coactivity pattern (inference-related pattern) detected in T2 session. The pattern is represented as a vector containing the contribution (weight) of each neuron’s spiking to the coactivity defining that pattern. Neurons with a weight above 2 s.d. of the mean were referred to as members (Red) (top). The temporal appearance of the pattern with the behaviour signature (just before the correct choice) (bottom). **g-i,** Activation strength (z-scored) of inference-related patterns (**g, i**), of stable training-related patterns (**h, i**). Hab, habituation session; NREM 1, NREM sleep after last habituation session; REM 1, REM sleep after last habituation session; NREM 2, NREM sleep after T1; REM 2, REM sleep after T1. Statistical comparisons were made using two-way repeated-measures analysis of variance (ANOVA) with Dunnett’s multiple comparison test (**g, h**). \**P* < 0.05; \*\**P* < 0.01; ns, not significant (*P* > 0.05). Data are presented as the mean ± standard error of the mean (s.e.m.).

Next, we employed combination of principal component and independent component analyses (PCA/ICA) to identify the neuronal assembly patterns and track their activity over time (Fig. 3f). We isolated different synchronized patterns during T2 session (Inference session) and tracked them over time. After that, we identified the inference-related patterns according to 3 criteria. First, they appeared in both T2 and T3 sessions just before the correct choice (entering and waiting in context B). Second, they appeared in T2 with an activation strength significantly higher than their strength during T1 session where the inferential behaviour was absent. Third, they did not appear during (B & C) and (C & D) sessions to confirm that these patterns are not representing context B preference or context D avoidance.

Upon investigating the temporal emergence of inference-related patterns, we found that they started to emerge from the randomization session and their activation strength increased gradually, peaking at REM sleep after T1 session (Fig. 3g, i). Strength of inference-related patterns was comparable between T2, T3 and REM 2 sessions, which was significantly higher than their strength in the earlier sessions. This result indicates that significant inference representations emerge during REM sleep after randomized training. Although inference-related patterns appeared during T1 session, their strength were not high enough to trigger inferential behaviour in T1. They were evident enough to elicit inferential behaviour in T2 and T3 sessions, suggesting that the level of pattern strength predict the task performance.

On the other hand, we extracted neuronal patterns representing the premise pairs during training sessions (see Methods). These patterns were divided into 3 groups; first, patterns activated significantly during initial training than during randomized training, named initial training-related patterns (Extended Data Fig. 7a). Second, patterns activated significantly during randomized training than during initial training, named randomized training-related patterns (Extended Data Fig. 7b). Third, patterns activated significantly during both initial and randomized training than during the earlier sessions, named stable training-related patterns (Stable “Tr, Rand”) (Fig. 3h). The stable “Tr, Rand” patterns representing both initial and randomized training were evident during REM sleep more than the initial training-related patterns, suggesting a higher contribution of stable “Tr, Rand” patterns in inference emergence (Extended Data Fig. 7c). Unlike inference-related patterns which reactivated during REM 2 more than NREM 2, stable “Tr, Rand” patterns reactivated similarly during NREM 2 and REM 2 sessions (Fig. 3h, i). Furthermore, comparing the activation strength of inference-related patterns and stable “Tr, Rand” patterns across sessions suggests that representations of learned and inferred information might be orthogonal (Fig. 3i).

### Coactivity code underlies the emergence of inferential behaviour representation

Understanding ACC computations during NREM 2 & REM 2 sleep sessions would explain the mechanism underlying the transitional increase of activation strength of inference-related patterns between NREM 2 and REM 2 (Fig. 3g), which predicts the inferential behaviour. To address this aim, we assessed the co-activation index between different patterns by calculating the number of pairwise synchronized Ca2+ activities within 200 ms between neurons constituting each pattern (Fig. 4a, b). Neurons contributing to the stable “Tr, Rand” patterns were co-activated significantly higher during NREM 2 than during REM 2 and awake sessions, representing a significant interaction (Fig. 4c). Furthermore, neurons belonging to inference-related patterns interacted with those belonging to stable “Tr, Rand” patterns significantly higher during REM 2 than during NREM 2 and awake sessions (Fig. 4c).

**Fig. 4.**
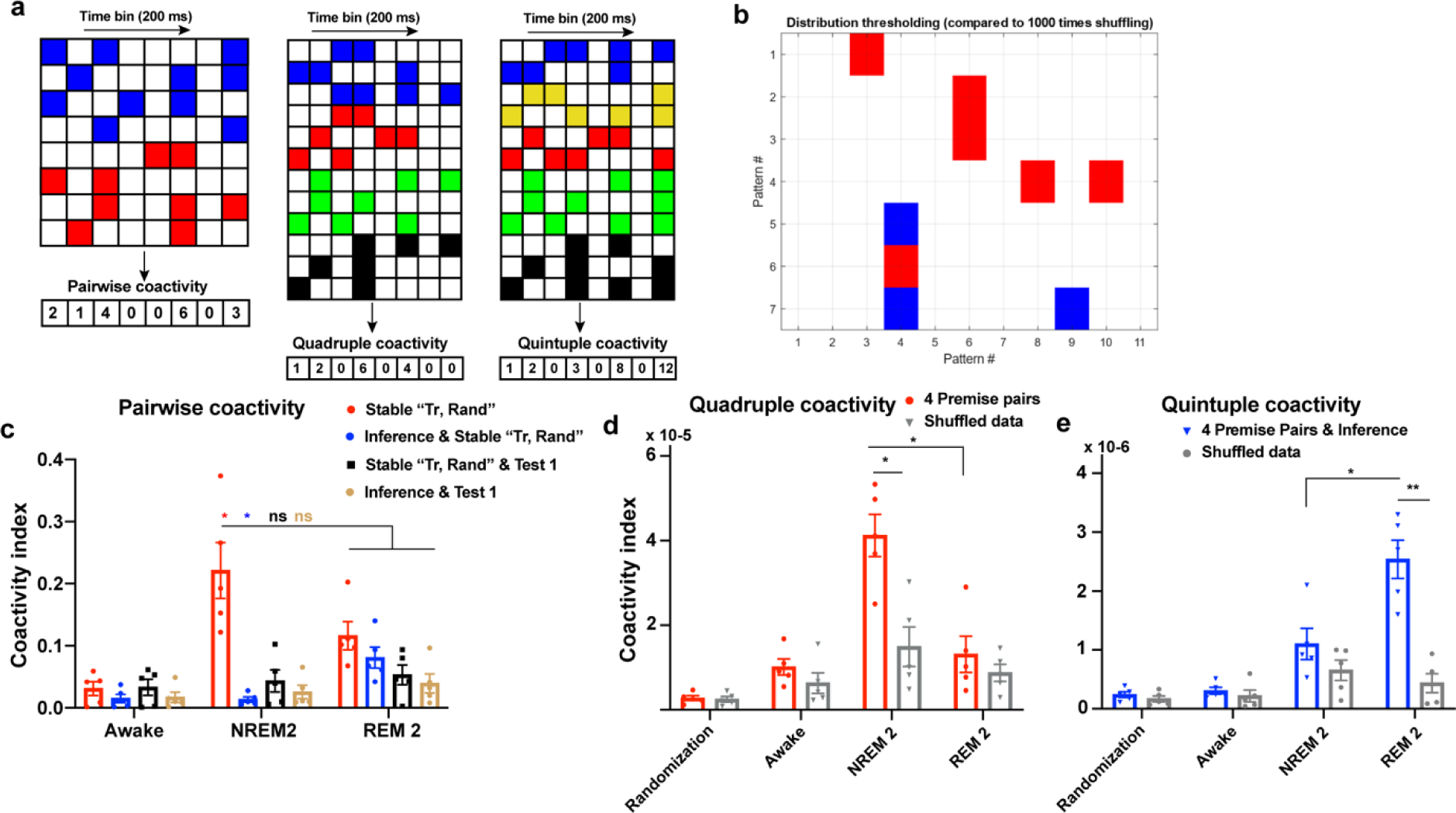
NREM and REM sleep coordinate to build synchronized inference representation with learned knowledge representation. **a,** Representative raster plots showing coactivity analysis. This analysis calculates coactivity by normalizing the number of synchronizations every 200 ms among different neuronal subpopulations representing different sessions. Coactivity analysis was done for neurons belonging to different patterns representing 2 sessions (Pairwise coactivity, left), representing 4 sessions (Quadruple coactivity, middle) and representing 5 sessions (Quintuple, right). **b**, Coactivity index between 2 different patterns compared to that appeared from shuffled data. Higher synchrony compared to shuffled data is represented with red color, while lower synchrony is represented with blue color. **c-e,** Coactivity index between 2 sessions (**c**), 4 sessions (**d**) and 5 sessions (**e**) with the shuffled data. Statistical comparisons were made using two-way repeated-measures analysis of variance (ANOVA) with Tukey’s multiple comparison test (**c-e**). \**P* < 0.05; \*\**P* < 0.01; ns, not significant (*P* > 0.05). Data are presented as the mean ± standard error of the mean (s.e.m.).

To examine the contribution of each premise pair (A&B, B&C, C&D, D&E) to the coactivity code, we extracted the patterns uniquely representing each premise pair and tracked the coactivity of their neurons. We counted the quadruple coactivation index between neurons representing the 4 individual premise pairs and found that they coactivated together during NREM 2 significantly more than during REM 2 session, which suggest organizing the learned information in hierarchy during NREM sleep (Fig. 4d). When we counted the quintuple coactivation index between neurons representing the 4 premise pairs and the inference patterns, we found higher synchronized activity during REM 2 than NREM 2 (Fig. 4e).

These results indicate the complementary role played by NREM and REM sleep. The learned knowledge is organized during NREM sleep, while it synchronizes with inferred information during REM sleep that may increase the strength of inference-related patterns leading to inferential behaviour. This two-step mechanism may explain the necessity of both NREM and REM sleep for building cognitive inference. Moreover, neurons constituting patterns of T1 session did not synchronize with those representing either training or inference sessions (Fig. 4c), supporting that T1 session is not critical for the inferential behaviour observed in T2 and T3.

### MEC→ACC crosstalk during REM sleep is sufficient to inspire inference from inadequate training

To establish a causal link between ACC coactivation code and inference processing, we sought to identify and manipulate a neural circuit that is involved in ACC coactivity formation. ACC coactivity code could rely on synaptic inputs from the connected upstream medial entorhinal cortex (MEC) which is thought to be involved in transitive inference^43,44^. We tested the sufficiency of such co-activation to elicit inferential behaviour from inadequate training. Mice were injected into MEC with AAV9 encoding light-sensitive neuronal activator (oChIEF) under the control of human synapsin (hSyn) 1 promoter (Fig. 5a). Mice were exposed to an incomplete/modified protocol of the transitive inference task, in which the randomized training, which is necessary to create inference (Fig. 1d, e), was replaced with optogenetic activation of axonal terminals of the MEC neurons in the ACC at 4 Hz during either NREM or REM sleep (Fig. 5b, c, and Extended Data Fig. 6b). All groups achieved the criterion for correct performance during training of the premises (Fig. 5d). Without both randomized training and MEC→ACC circuit activation, mice did not infer correctly, as demonstrated by a number of correct trials that was not significantly different from chance level in the B and D context tests (Fig. 5e). In contrast, artificial activation of the MEC→ACC network during REM, but not NREM, sleep resulted in inference in T2, T3, and T4, in which mice exhibited significantly more correct trials than those seen in T1, and significantly higher than the chance level; faster selection of the rewarded context was also found (Fig. 5e, f, and Extended Data Fig. 8). These results highlight the dissociable contribution of sleep stages (NREM and REM sleep) in inference processing, and indicates that activating the cortical network during REM sleep is sufficient for the creation of novel conclusions. All groups were proficient during testing of the non-inference and the premise pairs (Extended Data Fig. 2d).

**Fig. 5.**
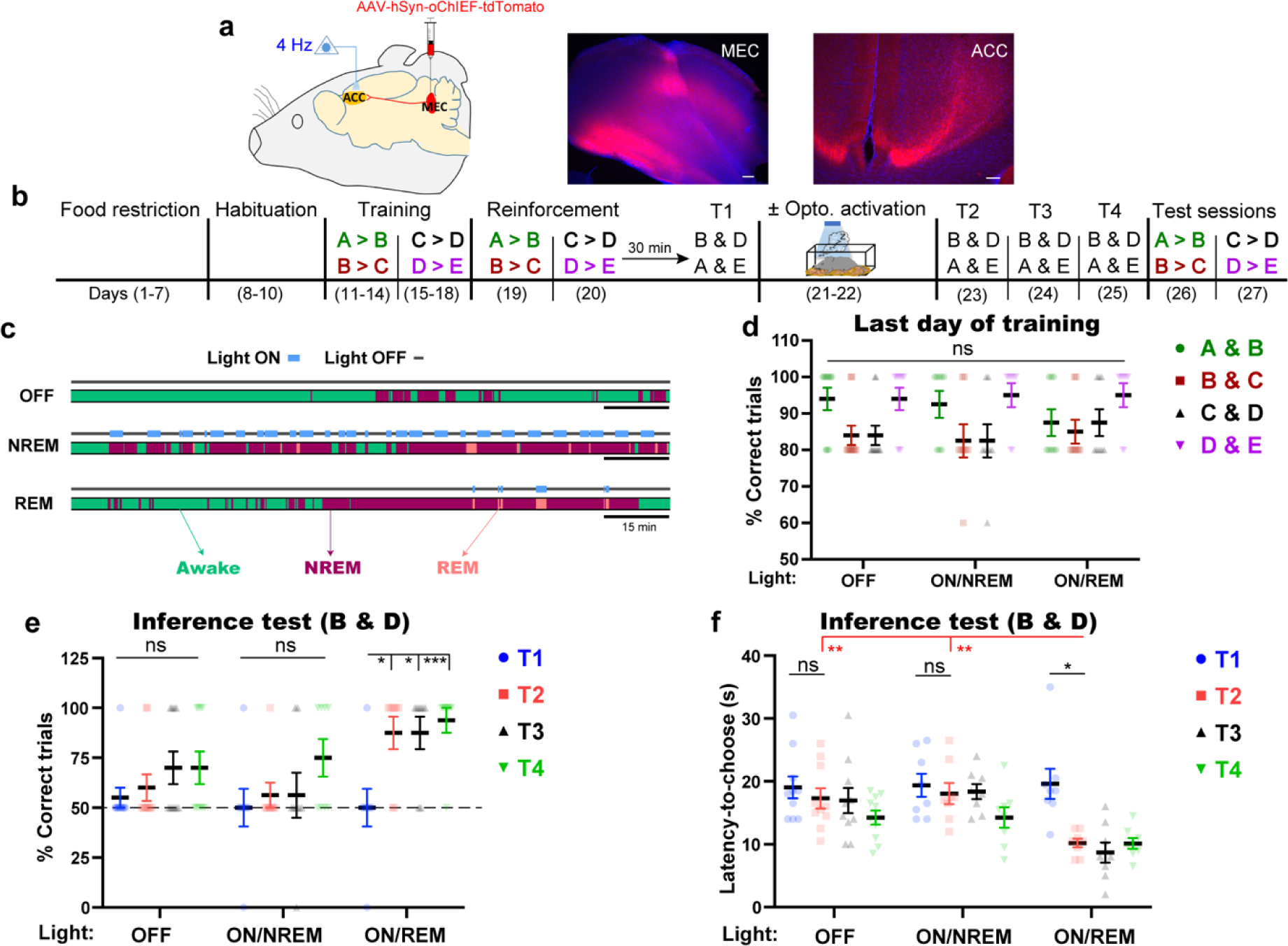
MEC→ACC crosstalk during only REM sleep is sufficient to inspire inference from inadequate training. **a,** The strategy for labelling MEC neurons and targeting their terminals in the ACC (left), and expression of oChIEF-tdTomato (red) in MEC neurons (middle) and their terminals in the ACC (right). Blue, 4′,6-diamidino-2-phenylindole (DAPI) staining. Scale bars, 100 µm. **b**, The behavioural schedule used to manipulate the MEC→ACC circuit during different sleep stages. **c**, Diagram showing the sleep stage-specific manipulation. Scale bar, 15 minutes. **d,** Performance during the last day of training (day 14 for A > B and B > C; day 18 for C > D and D > E) for each premise pair. **e-f,** Performance during test sessions; the percent of correct trials during the inference test (**e**), the latency time to choose during the inference test (**f**). T1, test 1; T2, test 2; T3, test 3; T4, test 4. *n* = 10 mice for the light off group; *n* = 8 mice for the non-rapid eye movement (NREM) sleep group; *n* = 8 mice for the rapid eye movement (REM) sleep group. Statistical comparisons were made using a two-way repeated-measures analysis of variance (ANOVA) (**d-f**) with Newman-Keuls multiple comparison test (**e**) and Tukey’s multiple comparison test (**d, f**). \**P* < 0.05; \*\**P* < 0.01; \*\*\*\**P* < 0.0001; ns, not significant (*P* > 0.05). Data are presented as the mean ± standard error of the mean (s.e.m.).

## Discussion

Our study clarifies the superiority of the subconscious brain over the conscious brain in restructuring existing stores of information before abstracting logical inference. During sleep, multiple related memories were connected to construct a knowledge structure, which facilitated innovative inference, a capacity that was not present shortly after learning. The discrepancy in the performance after awake and sleep manipulations indicates the necessity of offline, but not online, cortical activity to trigger inference. Sleep deprivation and optogenetic inhibition experiments revealed the ability to interfere with the emergence of inference while preserving memories of the original premises, thereby suggesting that stored knowledge and inferential information have unique representations and/or distinct locations in the brain. The artificial induction of inference by initiating communication between the MEC and ACC during REM sleep implies that the dynamic interplay between cortical neural codes for related experiences is critically important for the emergence of inferential behaviour.

Our study demonstrates that offline brain activity allows the abstraction of new information from previous experiences, which differs from previous findings showing the online emergence of inferential knowledge during subsequent experiences^38,45^. Furthermore, causal evidence for the link between inference and offline brain activity has been lacking^38,45^. Our study demonstrates the offline and spontaneous emergence of inference without prior priming, and provides causal evidence for the necessity and sufficiency of cortical activity during sleep to inspire inferential knowledge.

REM sleep is characterized by vivid dreams^26^ and could be initiated by activating specific dopaminergic neurons in amygdala^46^. Prior investigations into the role of REM sleep in cognition has found that it is necessary for memory consolidation^10^ and could be involved in creative problem solving by enhancing the formation of associative networks^23,26^. Here, we present evidence of a causal link between REM sleep and creativity using loss and gain of function experiments. Activating the MEC→ACC network at 4 Hz during REM sleep may provide an ideal setting for dreaming the inference, thus leading to innovative decisions. This suggestion is consistent with a previous study demonstrating that 4 Hz oscillations, dominant in PFC, were phase coupled with theta rhythm, leading to modulation of neuronal spikes in the PFC; hence supporting the processing of information^47^. Since theta rhythm (4-8 Hz) and network oscillations support transient synchronization between brain regions^47^, we hypothesize that artificial stimulation of MEC→ACC network at 4 Hz may mimic the physiological circuit mechanism of inference emergence.

The dissociable roles of REM and NREM sleep in processing the inference suggest that there are distinct offline mechanisms underlying the formation of knowledge networks. Since NREM sleep precedes REM sleep, we propose that NREM sleep prepares the knowledge network for REM-induced restructuring and the creation of novel associations^26^. This may explain why NREM sleep is crucial, but not sufficient, for the emergence of inference. The greater contribution of REM sleep over NREM sleep in inference development supports earlier findings that performance in anagram word puzzles is better after awakening from REM sleep than after awakening from NREM sleep^48^. Our Ca2+ imaging data suggests that NREM sleep may connect and organize the learned information in hierarchy, while REM sleep may build and stabilize the inference representation by creating new connections with the learned knowledge through co-reactivating their representations (Extended Data Fig. 9).

Ca2+ imaging unveiled a coactivity coding logic during offline state, which may link temporally segregated events and organize them in hierarchical order, allowing for building new information for future use. This demonstration is consistent with a previous study showing that synchronized offline reactivations are necessary for assimilating two subtly related memories, and eventually revealing the implicit commonality between them^42^. These data indicate that offline coactivation coding could underlie sleep-triggered creativity.

The capacity of REM sleep to facilitate inference from inadequate knowledge could inspire the development of novel approaches to boost cognitive performance. A recent study reported that dopamine signalling in amygdala is crucial for the transition from NREM to REM sleep^46^. Taken together, our study shed light on the mechanism of intellectual and cognitive impairments that occur in diseases that involve abnormalities in dopamine signalling like Parkinson’s disease and Attention-deficit hyperactivity disorder. The ability of REM sleep to flexibly restructure our knowledge could offer new behavioural repertoires to facilitate higher-order brain functions, such as implicit learning, decision making, and creative thinking.

## METHOD

### Animals

Naïve wild-type male C57BL/6J mice were purchased from Sankyo Labo Service Co. Inc. (Tokyo, Japan) and maintained on a 12 h light/dark cycle at a controlled temperature (24°C ± 3°C) and humidity (55% ± 5%) with free access to food and water. Mice used in behavioural experiments were 14–20 weeks old. All experimental procedures with animals were congruent with the guidelines of the National Institutes of Health. The Animal Care and Use Committee of the University of Toyama approved all animal experiments.

### Viral constructs

<u>For the optogenetic inhibition experiment (Fig. 2 and Extended Data Fig. 5)</u>, the AAV viral vector AAV-CaMKII-ArchT-eYFP (3.15 × 10^13^ vg/mL) was used. pAAV-CaMKII-ArchT-eYFP was kindly donated by Dr. K. Deisseroth. The recombinant AAV9 production was performed using a minimal purification method, and viral genomic titre was subsequently calculated as described previously^49^. Briefly, pAAV recombinant vector was produced using HEK293 T cells (AAV293; 240073, Agilent Tech, CA, USA) cultured in 15 cm dishes (Corning, NY, USA). Cultured cells were maintained in Dulbecco’s Modified Eagle Medium (D-MEM) (11995-065, GIBCO life technologies, USA) supplemented with 10% fetal bovine serum (FBS) (10270106, GIBCO life technologies, USA), 1% 2 mM L-Glutamine (25030-149, GIBCO Life Technologies, USA), 1% 10 mM non-essential amino acid (MEM NEAA 100x, 11140-050, GIBCO Life Technologies, USA), and 1% (100x) penicillin-streptomycin solution (15140-148, GIBCO Life Technologies, USA). Confluent (70%) HEK293 T cells were transfected using medium containing the constructed expression vector, pRep/Cap, and pHelper (240071, Agilent Technologies, Santa Clara, CA, USA) mixed with the transfection reagent polyethyleneimine hydrochloride (PEI Max, 24765-1, Polysciences, Inc., Warrington, PA, USA) at a 1:2 ratio (W/V). After 24 h, the transfection medium was discarded, and cells were incubated for another 5 days in an FBS-free maintenance medium. On day 6, the AAV-containing medium was collected and purified from cell debris using a 0.45 μm Millex-HV syringe filter (SLHV033RS, Merck Millipore, Germany). The filtered medium was concentrated and diluted with D-PBS (14190-144, GIBCO Life Technologies, USA) twice using the Vivaspin 20 column (VS2041, Sartorius, Germany) after blocking the column membrane with 1% bovine serum albumin (01862-87, Nacalai Tesque, Inc., Japan) in PBS. To further calculate the titre, degradation of any residual cDNA in the viral solution from production was first assured by benzonase nuclease treatment (70746, Merck Millipore, Germany). Subsequently, viral genomic DNA was obtained after digestion with proteinase K (162-22751, FUJIFILM Wako Pure Chemical, Osaka, Japan), extraction with phenol/chloroform/isoamyl alcohol 25:24:1 v/v, then precipitation with isopropanol and final dissolution in TE buffer (10 mM Tris [pH 8.0], 1 mM EDTA). Titre quantification for each viral solution, referenced to that of the corresponding expression plasmid, was done by real-time quantitative PCR (qPCR) using THUDERBIRD SYBR qPCR Master Mix (QRS-201, Toyobo Co., Ltd, Japan) with the primers 5′-GGAACCCCTAGTGATGGAGTT-3′ and 5′-CGGCCTCAGTGAGCGA-3′ targeting the inverted terminal repeat (ITR) sequence. The cycling parameters were adjusted as follows: initial denaturation at 95°C for 60 sec, followed by 40 cycles of 95°C for 15 sec and 60°C for 30 sec.

<u>For the *in vivo* Ca^2+^ imaging experiment (Fig. 3, 4)</u>, the AAV viral vector AAV-Syn-janelia-GCaMP7b (8.36 × 10^13^ vg/mL) was used. The pAAV-Syn-janelia-GCaMP7b^50^ was purchased from Addgene (Cambridge, MA, Plasmid #104489). The recombinant AAV9 production was performed using a minimal purification method, and viral genomic titre was subsequently calculated as described above and previously^49^.

<u>For the optogenetic activation experiment (Fig. 5)</u>, the AAV viral vector AAV-hSyn1-oChIEF-tdTomato (2.1 × 10^13^ vg/mL) was used. pAAV-hSyn1-oChIEF-tdTomato was purchased from Addgene (Cambridge, MA, Plasmid #50977). Recombinant AAV vectors were produced by transient transfection of HEK293 cells with the vector plasmid, an AAV3 rep and AAV9 vp expression plasmid, and an adenoviral helper plasmid, pHelper (Agilent Technologies, Santa Clara, CA), as previously described^51,52^.

### Surgery

Mice (10–14 weeks old) were given an intraperitoneal anaesthesia injection containing 0.75 mg/kg medetomidine (Domitor; Nippon Zenyaku Kogyo Co., Ltd., Japan), 4.0 mg/kg midazolam (Fuji Pharma Co., Ltd., Japan), and 5.0 mg/kg butorphanol (Vetorphale, Meiji Seika Pharma Co., Ltd., Japan) before being placed, when sedated, on a stereotactic apparatus (Narishige, Tokyo, Japan). After surgery, an intramuscular injection of 1.5 mg/kg atipamezole (Antisedan; Nippon Zenyaku Kogyo Co., Japan), an antagonist of medetomidine, was administered to boost recovery from sedation. Mice were home caged for 3 weeks to recover from surgery before initiating behavioural experiments. All virus injections were done using a 10 µL Hamilton syringe (80030, Hamilton, USA) that was fitted with a mineral oil-filled glass needle and wired to an automated motorized microinjector IMS-20 (Narishige, Japan).

<u>For the optogenetic inhibition experiments (Fig. 2 and Extended Data Fig. 5)</u>, 500 nL of AAV viral vector were injected at 100 nL min−1 bilaterally into either the ACC (from bregma: +0.3 mm anteroposterior [AP], ±0.35 mm mediolateral [ML]; from the skull surface: +1.5 mm dorsoventral [DV]) or prelimbic cortex (PL) (from bregma: +2.0 mm AP, ±0.35 mm ML; from the skull surface: +1.8 mm DV). The glass injection tip was maintained after injection at the target coordinates for an additional 5 min before being removed. Then, a double-guide cannula (C2002GS-5-0.7/SPC, diameter 0.29 mm, Plastics One Inc., USA) composed of two 0.7 mm-spaced stainless-steel pipes protruding for 2 mm from the plastic cannula body was bilaterally inserted either 1.0 mm ventral to the skull surface at the ACC coordinates or 1.3 mm ventral to the skull surface at the PL coordinates. Guide cannulas were fixed using dental cement (Provinice, Shofu Inc., Japan) that was used to fix micro-screws that were anchored into the skull near bregma and lambda. After complete fixation, a dummy cannula (C2002DCS-5-0.7/SPC, protrusion 0 mm, Plastics One Inc., USA) was attached to the guide cannula to protect from particulate matter.

In parallel, a custom-built electroencephalogram/electromyography (EEG/EMG) 5-pin system was installed into the skull as previously described^40^. Briefly, electrodes were screwed into the skull over the parietal cortex for EEG recording, over the right cerebellar cortex as a ground, and over the left cerebellar cortex as a reference. Additionally, two wires were implanted into the neck muscle for EMG recording. Finally, dental cement was used to fix all system screws in place.

<u>For the optogenetic activation experiment (Fig. 5)</u>, 500 nL of the AAV viral vector was injected at 100 nL min−1 bilaterally into the MEC (from bregma: +4.8 mm AP, ±3.3 mm ML, +3.3 mm DV). The remaining procedure was as described earlier in the optogenetic inhibition section.

<u>For the Ca^2+^ imaging experiment (Fig. 3, 4)</u>, 500 nL of AAV_9_-CaMKII::G-CaMP7 was injected at 100 nL min^−1^ unilaterally into the left ACC (+1.0 mm AP, +0.35 mm ML, +1.5 mm DV). After 2 weeks of recovery from AAV injection surgery, re-anesthetized mice were placed once again on a stereotactic apparatus to implant a gradient index (GRIN) lens^8,42,53^ (0.5 mm diameter, 4 mm length; Inscopix Inc., USA) into the centre of injection (from the skull surface: +1.2 mm DV) using custom-made forceps attached to a manipulator (Narishige, Japan). A low-temperature cautery was used to emulsify bone wax into the gaps between the GRIN lens and the skull, and then the lens was anchored in place using dental cement. Additionally, a custom-built EEG/EMG 5-pin system was installed and cemented into the skull, as mentioned earlier. Three weeks after GRIN implantation, mice were re-anesthetized and placed back onto the stereotactic apparatus to set a baseplate (Inscopix Inc.), as described previously^42,53^. In brief, a Gripper (Inscopix Inc.) holding a baseplate attached to a miniature microscope (nVista HD v3; Inscopix, Inc.) was lowered over the previously set GRIN lens until visualization of clear vasculature was possible, indicating the optimum focal plane. Dental cement was then applied to fix the baseplate in position to preserve the optimal focal plane. Mice recovered from surgery in their home cages for 1 week before behavioural imaging experiments began.

### Transitive inference task

The behavioural experiments were done in soundproof room. The arena consists of 5 different contexts (A, B, C, D, and E; Fig. 1a). Context A was a grey triangle (200 mm length × 300 mm height) with a smooth, grey acrylic floor and walls without any patterns. Context B was a grey square (200 mm length × 200 mm width × 300 mm height) with a green plastic floor with a pointy texture and walls without any patterns. Context C was a blue square (200 mm length × 200 mm width × 300 mm height) with a smooth, blue acrylic floor and walls without patterns. Context D was a transparent square (200 mm length × 200 mm width × 300 mm height) with a spongy, orange floor and walls that were covered with a characteristic pattern (black circles on a white background). Context E was a transparent triangle (200 mm length × 300 mm height) with a smooth, transparent acrylic floor and walls that were covered with a characteristic pattern (black vertical lines on a white background).

#### Food restriction

Mice were kept under a food restriction protocol during the task. Mice were provided with one 3 g food pellet and one 0.05 g sucrose tablet per day until mice reach 80–85% of their original weight. The sucrose tablets were put in a small Petri dish inside the home cage during the first two stages (food restriction and arena habituation); the dish was removed after starting the training stage and was never put it back, which served to teach mice that they would no longer receive any sucrose tablets in the home cage and the only way to receive the reward in the contexts was to perform the task correctly.

#### Arena habituation

On the first day, mice were put in each context separately for 5 min. The contexts were closed to force mice to explore each context for the entire 5 min. The exposure to the contexts was random across mice (for example, one mouse was placed in A then C then D then B then E, and another mouse was placed in C then B then E then A then D). On the second and third days, all contexts were open and mice were put in the start point for free exploration of the whole arena for 15 min.

#### Training

The 14-day training phase was divided into 4 consecutive days for the first two pairs, 4 days for the other two pairs, 2 days reinforcement, and 4 days randomization. Every day, mice were exposed to two sessions, and each session consisted of five trials for a premise pair without inter-trial intervals. The four premise pairs were A > B, B > C, C > D, and D > E. Therefore, the hierarchy was A > B > C > D > E. For the incomplete hierarchy experiment (Extended Data Fig. 4), the four premise pairs were A > B, B > C, E > D, and D > C. Therefore, the hierarchies were A > B > C and E > D > C. Premise pair training was considered successful when mice reached 80% correctness.

In the first trial of every session, mice were put at the starting point and allowed to explore the arena and enter the two contexts to identify the learned pair. After entering both contexts once, if they entered the correct context again and remained there for more than 10 sec, they received the sucrose tablet, and the trial ended after the tablet had been consumed. If they stayed in the non-rewarded context (wrong choice) for more than 10 sec, the trial ended, and in the next correct trial they received the sucrose tablet after 20 sec (rather than 10 sec). After each trial, mice were directly returned to the starting point to start the next trial without any inter-trial interval. After completing all trials, mice were returned to their home cage for 30 min before starting the next session. After finishing two sessions, mice returned to their home cages and received a 2-g food pellet.

For randomization during training, mice were exposed to new combinations of premise pairs everyday for 4 days. These combinations were different from those presented during the initial training and reinforcement. For example, (A & B) pair was co-presented with (B & C) pair on the same day during the initial training and reinforcement, while during randomization stage (A & B) pair was co-presented with (C & D) and (D & E) pairs on two different days; and the same was applied on the other pairs to ensure the complete interaction between all premise pairs. The order of presenting the premise pairs is different across animals within the same group (Table 1). This is different from training with reinforcement, in which the (A & B) pair is usually presented with the (B & C) pair, and the (C & D) pair is usually presented with the (D & E) pair on the same day.

**Table 1:** Example of the order of presenting the premise pairs during randomization stage.

| Randomization Day # | 1 | 2 | 3 | 4 |
| --- | --- | --- | --- | --- |
| Day # in the protocol | Day # 11 | Day # 12 | Day # 13 | Day # 14 |
| Mouse # 1 | A > B<br>C > D | B > C<br>C > D | A > B<br>D > E | B > C<br>D > E |
| Mouse # 2 | A > B<br>D > E | A > B<br>C > D | B > C<br>D > E | B > C<br>C > D |
| Mouse # 3 | B > C<br>C > D | B > C<br>D > E | A > B<br>D > E | A > B<br>C > D |

<u>For the optogenetic activation experiment (Fig. 5 and Extended Data Fig. 8),</u> training was done for 10 days only without the 4 days of randomization.

#### Testing

After the training phase, mice were exposed to two types of testing for 4 consecutive days. The inference test utilized a novel pair (B & D), while the non-inference test utilized another novel pair (A & E). Test 1 was done on the last day of training, while tests 2, 3, and 4 were done on the following 3 days.

Every day, mice were exposed to two sessions, each of which consisted of two trials for each test without any inter-trial intervals. After completing all trials, mice were returned to the home cage for 30 min before starting the next session. The protocol of test sessions was the same as for the training sessions. When they made the correct choice, mice were rewarded with a sucrose tablet in context B, except for the inference group (Fig. 1f) and both awake and sleep groups (Extended Data Fig. 5). Both the B and D contexts were square and located at the end of the arena (the same distance from the starting point) to avoid any preference that was due to geometry and distance from the starting point. Performance during the B and D test was also not due to right or left training preferences, because the design of the premise pairs was based on an equal distribution of right and left preferences (the correct choice was located to the left in two pairs and to the right in the other two pairs). After completing the inference and non-inference tests, mice were tested with the original premise pairs to confirm that the performance during the inference test was due to remembering the original memories. The arena was cleaned using water and 70% ethanol after each subject. Sleep deprivation (Fig. 1) was done for 4 h by gentle touching of the home cage after test sessions (after T1, T2, T3, and T4).

<u>For the optogenetic inhibition experiments (Fig. 2 and Extended Data Fig. 5)</u>, optogenetic inhibition of the ACC or PL was induced during wakefulness or sleep in a 4 h session. On each testing day, immediately after the test sessions, mice were anesthetized using isoflurane and two branch-type optical fibres (internal diameter, 0.25 mm) were inserted and fitted into their housing with a cap, which anchored the inserted optical fibres with screws around the guide cannula. The tip of the optical fibre protruded 0.2 mm below the guide cannula (ACC, DV 1.2 mm from the skull surface; PL, DV 1.5 mm from the skull surface). Mice attached to the optic fibres were then placed in a sleep box and simultaneously connected to an EEG/EMG recording unit and an optical swivel wired to a laser unit (9–12 mW, 589 nm). The delivery of continuous light was manually controlled. After the session, mice were detached from the EEG/EMG recording and light delivery systems and the fibres were removed from the cannula under anaesthesia.

<u>For the *in vivo* Ca^2+^ imaging experiment (Fig. 3, 4)</u>, attachment and removal of the microendoscope was performed under 3% isoflurane anesthesia. Mice attached to the microendoscope were returned to their home cages to recover for at least 30 min before and after the behavioral session. Three days before context habituation sessions, mice were habituated to the microendoscope attachment for 10 min per day in their home cages. Calcium imaging data acquisition started from the last day of context habituation. Mice were exposed to the above-mentioned behavioural protocol with slight modification in the initial training stage. Mice were exposed to two particular premise pairs for 2 successive days, then they encountered a single premise pair per day on the subsequent 2 days to collect the corresponding Ca2+ data of each premise pair on a separate day. Offline Ca2+ imaging data was collected from the sleep period (NREM 1 & REM 1) after contexts habituation, the awake session after Test 1 (Day 24) and the sleep period (NREM 2 & REM 2) after Test 1. Ca2+ imaging data was collected from the first 2 min of NREM sleep (single NREM epoch that lasts for more than 2 min) and from the first 2 min of REM sleep (multiple REM epochs; number of epochs across animals is 3±1).

<u>For the optogenetic activation experiment (Fig. 5)</u>, optogenetic activation to MEC terminals in the ACC was performed during NREM or REM sleep for two consecutive days. One day after test 1, mice were anesthetized using isoflurane and two branch-type optical fibres (internal diameter, 0.25 mm) were inserted and fitted into their housing with a cap, which anchored the inserted optical fibres by screws around the guide cannula. The tip of the optical fibre protruded 0.2 mm below the guide cannula (DV 1.2 mm from the skull surface). Mice attached to the optic fibres were then placed in a sleep box, and simultaneously connected to an EEG/EMG recording unit and an optical swivel wired to a laser unit (9–12 mW, 473 nm). The delivery of laser stimulation (4 Hz, 15 msec pulse width) was manually controlled using a schedule stimulator in time-lapse mode (Master-8 pulse stimulator, A.M.P.I.). In REM-stimulated mice, 4 Hz light stimulation was delivered during all REM sleep that occurred within the 4 h sleep session; the total duration of light stimulation was less than 7 min per mouse. For NREM-stimulated mice, light stimulation was delivered for a maximum of 3 min per epoch with a 3 min inter-epoch interval. After the session, the mice were detached from the EEG/EMG recording and light delivery systems and the fibres were removed from the cannula under anaesthesia.

<u>For the optogenetic activation experiment (Extended Data Fig. 8)</u>, the procedure was the same as that described in the previous paragraph, related to Fig. 5, with a modification in the stimulation protocol; the total duration of light stimulation was modified to be less than 7 min per mouse, with a maximum of 3 min per epoch. This modification was done to avoid prolonged light stimulation and to mimic the protocol used in REM-stimulated animals.

### Behavioural analysis

All sessions were captured with an overhead web camera (Logitech HD pro C920) mounted on a vertical stand. The time spent in each context during the habituation phase was counted. During training and testing, the trial was considered to be correct if mice spent > 10 sec in the assigned context. The percent of correct trials during training and testing sessions was calculated as follows: % correct trials = (number of correct trials / total number of trials) * 100. We set 80% correct trials during the training phase as the criterion for successful learning of the premise pairs.

### *In vivo* Ca^2+^ imaging data acquisition and analyses

Ca^2+^ signals produced from G-CaMP7 protein were captured at 20 Hz with nVista acquisition software (Inscopix, CA, USA) at the maximum gain and optimal power of LED of nVistaHD. Ca^2+^ imaging movie recordings of all behavioural sessions were then extracted from nVista Data acquisition (DAQ) box (Inscopix, CA, USA). Using Inscopix data processing software (Inscopix), movies were temporally stitched together to create a full movie that contained recordings of all sessions across days, which were spatially down-sampled (2x), and then corrected for across-session motion artefacts against a reference frame that was chosen from any session that had a clear landmark “vasculature”. Further motion correction was then applied using Inscopix Mosaic software, as previously described^8^. The full movie was then temporally divided into the individual sessions using Inscopix Mosaic software. Each movie of individual sessions was then low bandpass filtered using Fiji software (NIH) to reduce noise, as described previously^8^. The fluorescence signal intensity change (ΔF/F) for each session was subsequently calculated using Inscopix Mosaic software according to the formula ΔF/F = (F – Fm)/Fm, where F represents each frame’s fluorescence and Fm is the mean fluorescence for the whole session’s movie. Afterwards, movies representing each session were re-concatenated again to generate the full movie for all sessions in the ΔF/F format. Finally, cells were identified using an automatic sorting system, HOTARU, and each cell’s Ca^2+^ signals over time were extracted in a (Ď; time × neuron) matrix format, as previously described^8^. Further processing was performed using codes written in MATLAB to remove the low frequency fluctuation and background noise by subjecting output Ca^2+^ signals to high-pass filtering with a 0.01 Hz threshold, and to then calculate z-scores from the mean of each session, whereby negative values were replaced with zero. Ca^2+^ events were finally extracted after cutting off signals below 3 SD from the local maxima of the ΔF/F signal of each session.

### Neural ensembles analyses

Neural ensembles representing a range of neurons co-firing together was calculated using unsupervised statistical method using PCA-ICA analysis as previously described^54,55^. Initially, frames were binned to 0.2 s, then to prevent the bias due to differences in the average firing rates, neuronal Ca^2+^ transients were normalized for each neuron by a ***z***-score transform:

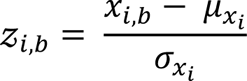

where for neuron *i*, *z*_i,b_ is its ***z***-scored Ca^2+^ transients in bin *b*, *x*_*i*,*b*_ its Ca^2+^ transients in bin *b*, *μ_x_i__* its mean Ca^2+^ transients across all bins and *σ_x_i__* the s.d. of its Ca^2+^ transients. The calculation of the principal components and the determination of the number of significant components were carried out according to previously presented methods.

To quantify the extent to which ACC neurons contribute to the formation of neuronal ensembles, we applied reconstruction ICA to significant principal components as described previously^54,55^. Ensemble activation strength was then calculated through:

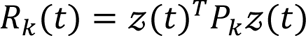

where *R*_*k*_(*t*), the activation strength of ensemble *k* at time *t*

Wilcoxon rank-sum test was then used to reveal which of the identified ensembles are significantly different between reference and target sessions. *R*_*k*_(*t*) > 2 were set as a threshold for ensembles to be included in the test.

### Coactivity index analysis

The coactivity index was calculated as described previously^53^. The z-scores of neuronal Ca2+ events were calculated as described above. The z-scores were binarized by thresholding (> 3 Standard Deviations from the ΔF/F signal) at the local maxima of the ΔF/F signal, and then were temporally down-sampled from 20 to 5 Hz data (200 ms binning). Subsequently, the coactivation index consisting of the number of synchronized activities among neurons - constituting different patterns from different sessions-in each 200 ms time window, was calculated and normalized in each behavioural session. The following equations are used for the calculations of pairwise, quadruple and quintuple coactivity index:

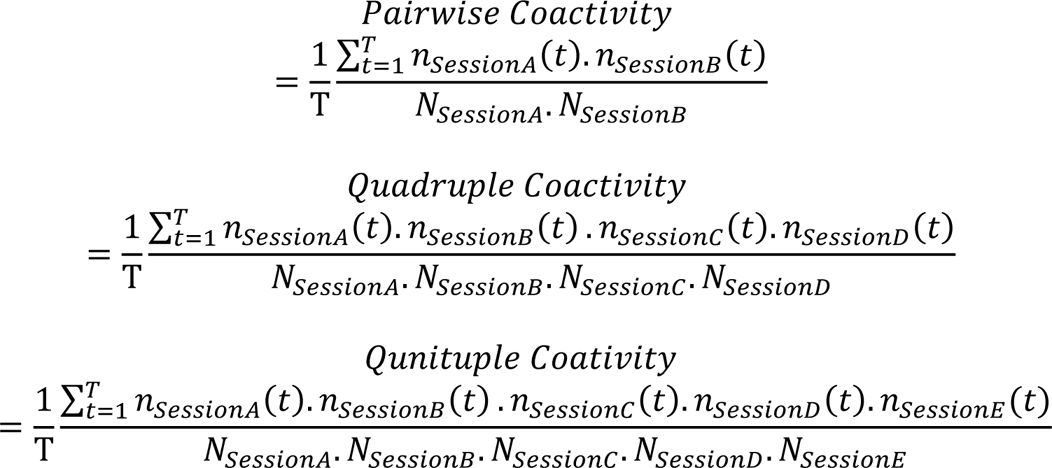

Where A, B, C, D & E are different behavioral sessions; *n_SessionA_*(*t*) is the number of cells, constituting significant patterns in Session A, that were active in the time bin t; *N_SessionA_* is the total number of Session A cells; and T is the total number of time bins for each session.

To exclude the possibility that the observed coactivity results from the difference in activities among cells and sessions, we generated shuffled data (1,000 surrogate values) by using the “circshift function of MATLAB”^56^.

### Sleep detection data acquisition and online state detection

All EEG/EMG recordings were performed using OpenEx Software Suite (Tucker Davis Technologies, USA), as previously described^40^ with minor modifications. Briefly, EEG signals were amplified and filtered at 1–40 Hz, while 65–150 Hz was used for EMG; signals were then digitized at a sampling rate of 508.6 Hz. Sleep stages were differentiated using an algorithm file that enabled the calculation and analysis of the EMG root mean square value (RMS), EEG delta power (1–4 Hz) RMS, and EEG theta power (6–9 Hz) RMS. The EMG RMS threshold was optimized according to each subject. Mice was judged to be awake when the EMG RMS exceeded the set threshold value and remained unchanged for three successive 3 sec checkpoints. However, when EMG RMS was lower than the threshold, sleep stage differentiation was concluded based on the delta/theta (d/t) ratio value. Briefly, if the d/t ratio exceeded 1 for the consecutive 9 sec checking period, the stage was classified as NREM sleep; conversely, it was classified as REM sleep when it was less than 1 for a consecutive 9 sec. The state classified by the program was also confirmed by the experimenter through visual inspection of mouse activity and EEG delta-dominant (0.5–4 Hz) or theta-dominant (4–9 Hz) waveforms. EEG/EMG traces recorded during sleep sessions were then extracted using MATLAB codes.

### Histology

After the optogenetic manipulation experiments, mice were deeply anesthetized with 1 mL of combination anaesthesia and perfused transcardially with PBS (pH 7.4) followed by 4% paraformaldehyde in PBS. The brains were extracted, then further immersed in paraformaldehyde in PBS for 12–18 h at 4°C. Subsequently, fixed brains were mixed with 25% sucrose in PBS for 36–48 h at 4°C before final storage at –80°C. To obtain coronal sections, brains were sliced into 50 μm sections using a cryostat (Leica CM3050, Leica Biosystems) and were then washed in PBS-containing 12-well culture plates (Corning, NY, USA). The sections were further incubated at room temperature for 1 h with a blocking buffer (3% normal donkey serum; S30, Chemicon by EMD Millipore, Billerica, MA, USA) in PBS solution containing 0.2% Triton X-100 and 0.05% Tween 20 (PBST). After incubation, the buffer was discarded and rat anti-GFP (04404-84, GF090R, Nacalai Tesque Inc., Japan) primary antibody (1:500) in blocking solution was added for further incubation at 4°C for 24–36 h. At the end of the incubation period, the primary antibody was removed and sections were washed with 0.2% PBST three times for 10 min each. After washing, sections were treated with a complementary secondary antibody, (1:1000) donkey anti-rat IgG Alexa Fluor 488 (A21208, Molecular Probes) in blocking buffer solution at room temperature for 2–3 h. Finally, the sections were treated with DAPI (1 μg/mL, Roche Diagnostics, 10236276001) for nuclear staining and washed with 0.2% PBST three times for 10 min each before being mounted onto glass slides with ProLong Gold antifade reagent (Invitrogen).

### Confocal microscopy

Images were acquired using a confocal microscope (Zeiss LSM 780, Carl Zeiss, Jena, Germany) with 20× Plan-Apochromat objective lens. All parameters, such as the photomultiplier tube assignments, pinhole sizes, and contrast values, were standardized within each magnification and experimental condition.

### Statistics

Statistical analyses were performed using Prism 8 (GraphPad Software, San Diego, CA, USA). Multiple-group comparisons were performed using an ANOVA with post hoc tests, as shown in the corresponding Fig. legends. Quantitative data are presented as the mean ± SEM.

## Data and Code Availability

All data, codes, and resources that supported the findings of this study are available upon reasonable request.

## Acknowledgments

We thank Khaled Ghandour and Ali Choucry (University of Toyama) for their valuable discussions and suggestions. We thank all members of the Inokuchi Laboratory for their discussions and support. We thank Mika Ito and Naomi Takino (Jichi Medical University) for their help with the production of the AAV vectors. Also, we thank Noriaki Ohkawa (Dokyyo University) for his help in providing the materials used to build the behavioural arena. We would like to thank Karl Deisseroth (Stanford University) for providing us with pAAV-CaMKII-ArchT 3.0-EYFP. We also thank Ayumu Konno and Hirokazu Hirai (Gunma University) for providing us with the virus preparation protocol.

This work was supported by the JSPS KAKENHI (grant number JP18H05213, JP23H05476), the Core Research for Evolutional Science and Technology (CREST) program (JPMJCR13W1) of the Japan Science and Technology Agency (JST), a Grant-in-Aid for Scientific Research on Innovative Areas “Memory dynamism” (JP25115002) from MEXT, and the Takeda Science Foundation to K.I.; JSPS KAKENHI (grant number JP19K16892) to K.A.; and the Uehara Memorial Foundation scholarship to M.A.; JST SPRING scholarship (grant number JPMJSP2145) to A.I.

## Author contributions

K.A. and K.I. designed the experiments and wrote the manuscript. K.A., K.C., M.A. and A.I. performed the experiments. K.A. analysed the data. S.M. and R.O-S. prepared adeno-associated viruses. K.I. supervised the entire project.

---

**Extended Data Fig. 1.**
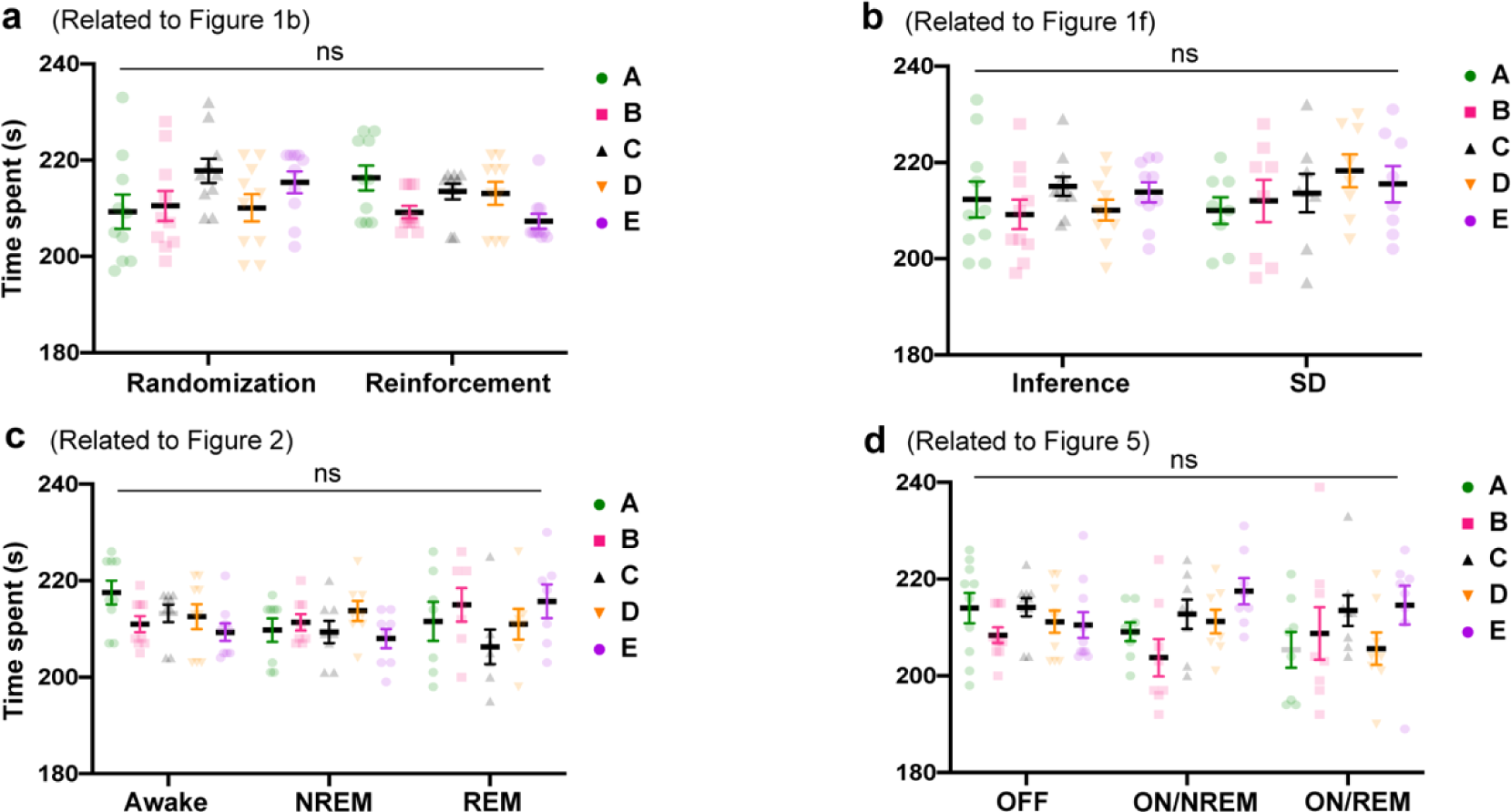
Mice had no innate preference to any context during habituation. **a-d,** Time spent in each context during the habituation phase. The number of animals in each panel is mentioned in the corresponding Figure legend. SD, sleep deprivation; NREM, non-rapid eye movement; REM, rapid eye movement. Statistical comparisons were made using a two-way repeated-measures analysis of variance (ANOVA) with Tukey’s multiple comparison test (**a-d**). ns, not significant (*P* > 0.05). Data are presented as the mean ± standard error of the mean (s.e.m.).

**Extended Data Fig. 2.**
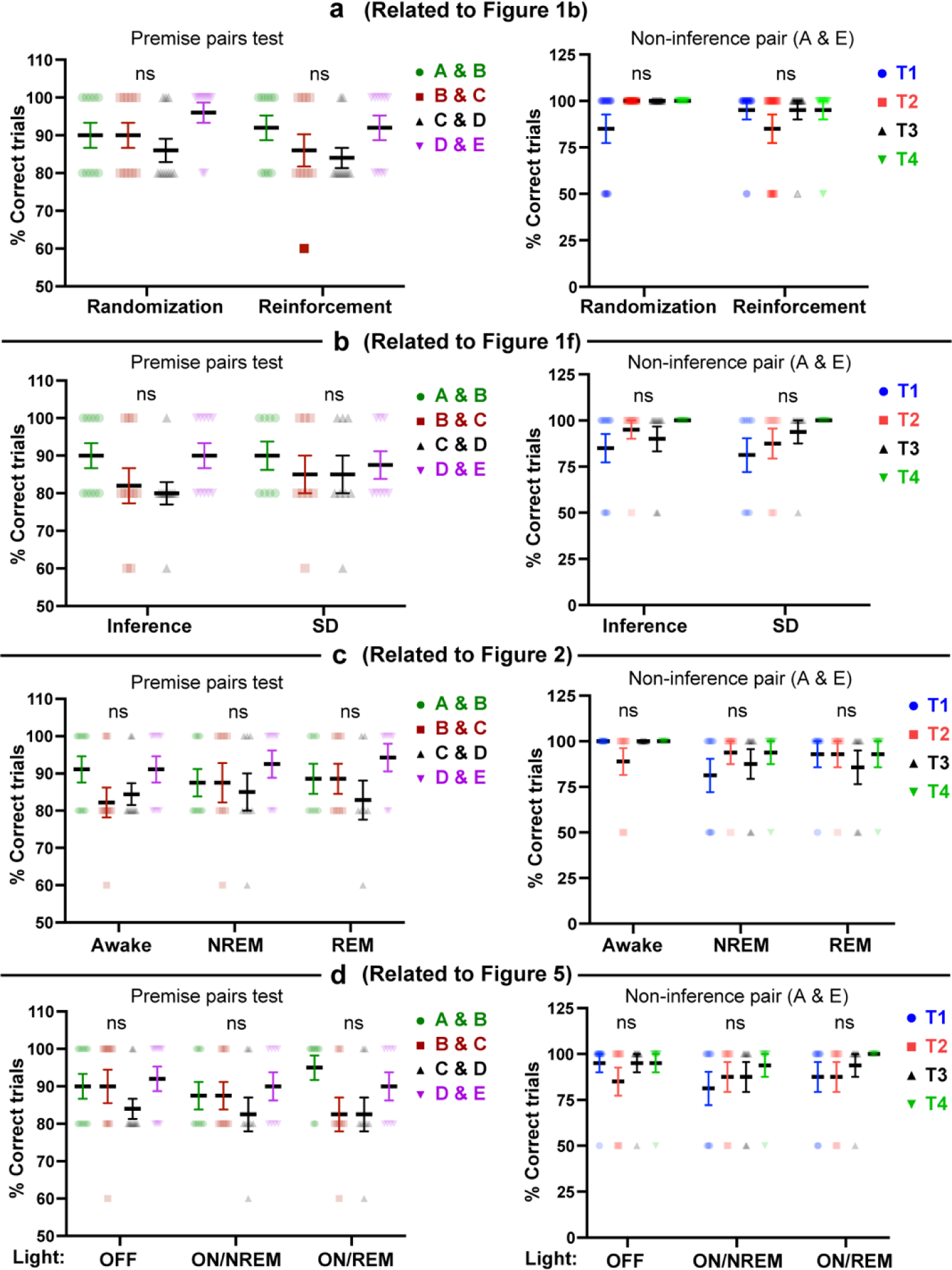
Correct performance during testing original premise pairs and non-inference pair. **a-d,** Performance during test sessions; the percent of correct trials during the original premise pairs test (left) and during the non-inference test (right). T1, test 1; T2, test 2; T3, test 3; T4, test 4. SD, sleep deprivation; NREM, non-rapid eye movement; REM, rapid eye movement. The number of animals in each panel is mentioned in the corresponding Figure legend. Statistical comparisons were made using a two-way repeated-measures analysis of variance (ANOVA) with Tukey’s multiple comparison test (**a**-**d**). ns, not significant (*P* > 0.05). Data are presented as the mean ± standard error of the mean (s.e.m.).

**Extended Data Fig. 3.**
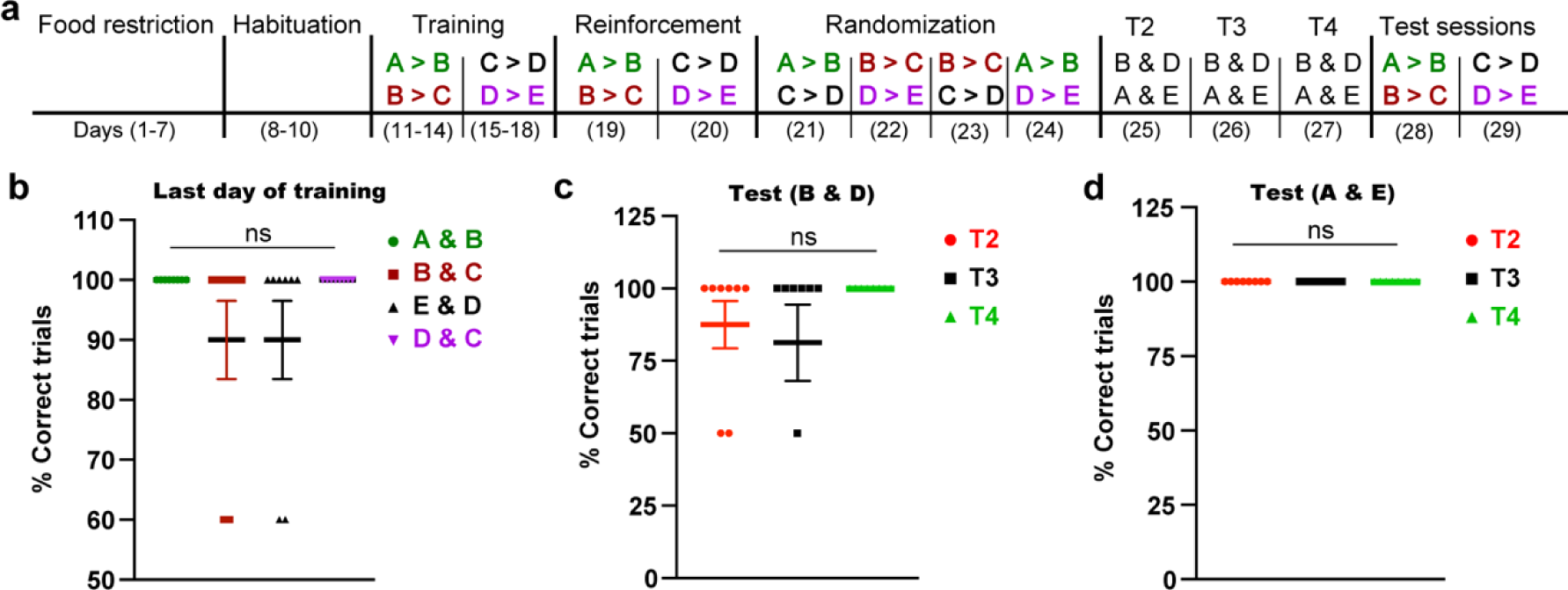
Inferential behaviour does not require priming by preexposure to the testing environment. **a,** The behavioural schedule used to test the impact of the complete hierarchy on inference development. The order of presentation of the premise pairs during the randomization stage is different across animals within the same group (See Methods; Table 1). **b**, The performance during the last day of training (day 14 for A > B and B > C; day 18 for E > D and D > C) for each premise pair was calculated as the percent of correct trials out of the total number of trials. **c-d,** Performance during test sessions; the percent of correct trials during the inference test (**c**) and during the non-inference test (**d**). Statistical comparisons were made using a one-way repeated-measures analysis of variance (ANOVA) with Tukey’s multiple comparison test; n=8 mice; ns, not significant (*P* > 0.05). Data are presented as the mean ± standard error of the mean (s.e.m.).

**Extended Data Fig. 4.**
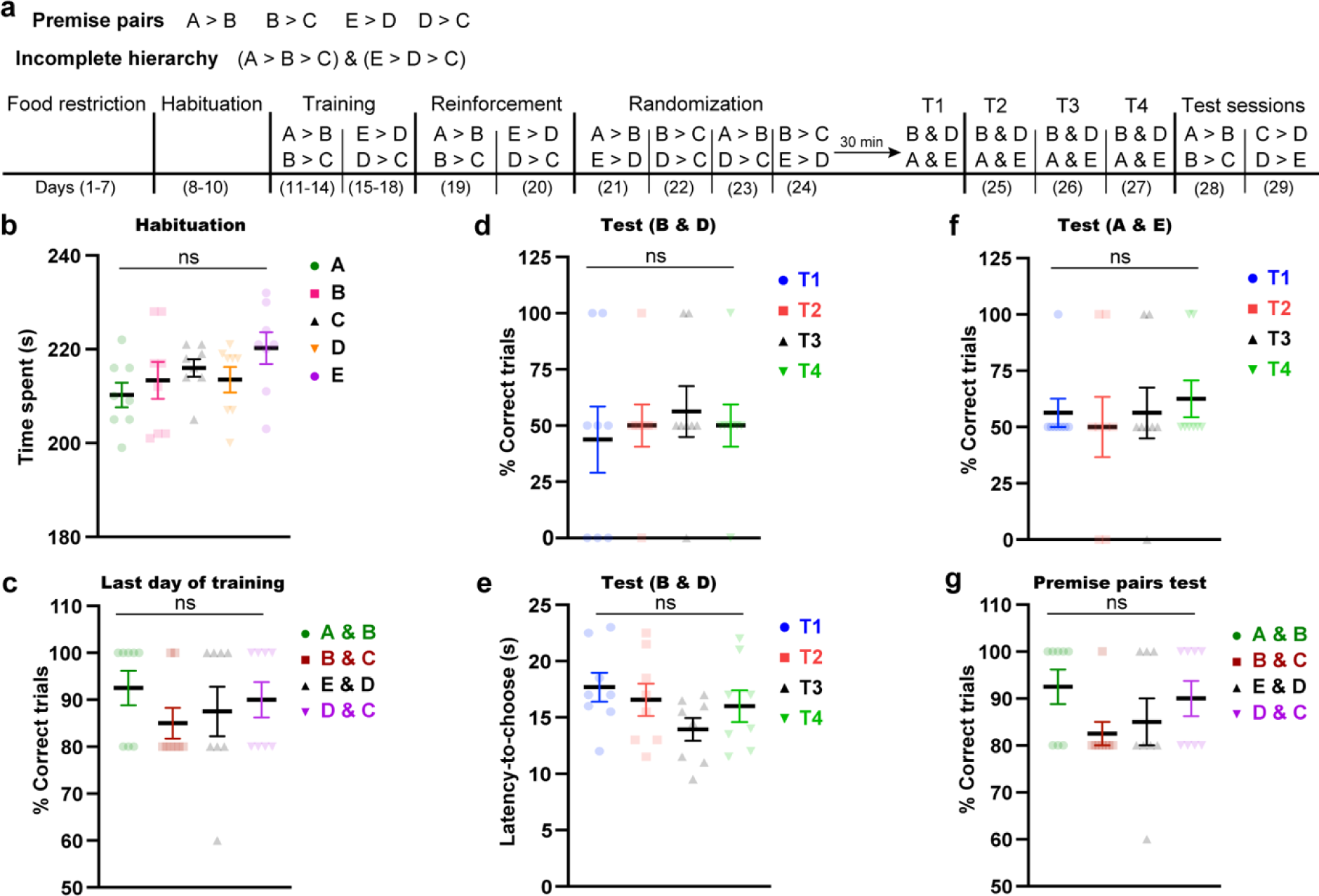
Without the complete hierarchy, mice could not infer correctly. **a,** The behavioural schedule used to test the impact of the complete hierarchy on inference development. **b**, Time spent in each context during the habituation phase. **c,** The performance during the last day of training (day 14 for A > B and B > C; day 18 for E > D and D > C) for each premise pair was calculated as the percent of correct trials out of the total number of trials. **d-g,** Performance during test sessions; the percent of correct trials during the inference test (**d**), the latency time to choose during the inference test (**e**), during the non-inference test (**f**), during the original premise pairs test (**g**). T1, test 1; T2, test 2; T3, test 3; T4, test 4. *n* = 8 mice. Statistical comparisons were made using a one-way repeated-measures analysis of variance (ANOVA) with Tukey’s multiple comparison test (**b**-**g**). ns, not significant (*P* > 0.05). Data are presented as the mean ± standard error of the mean (s.e.m.).

**Extended Data Fig. 5.**
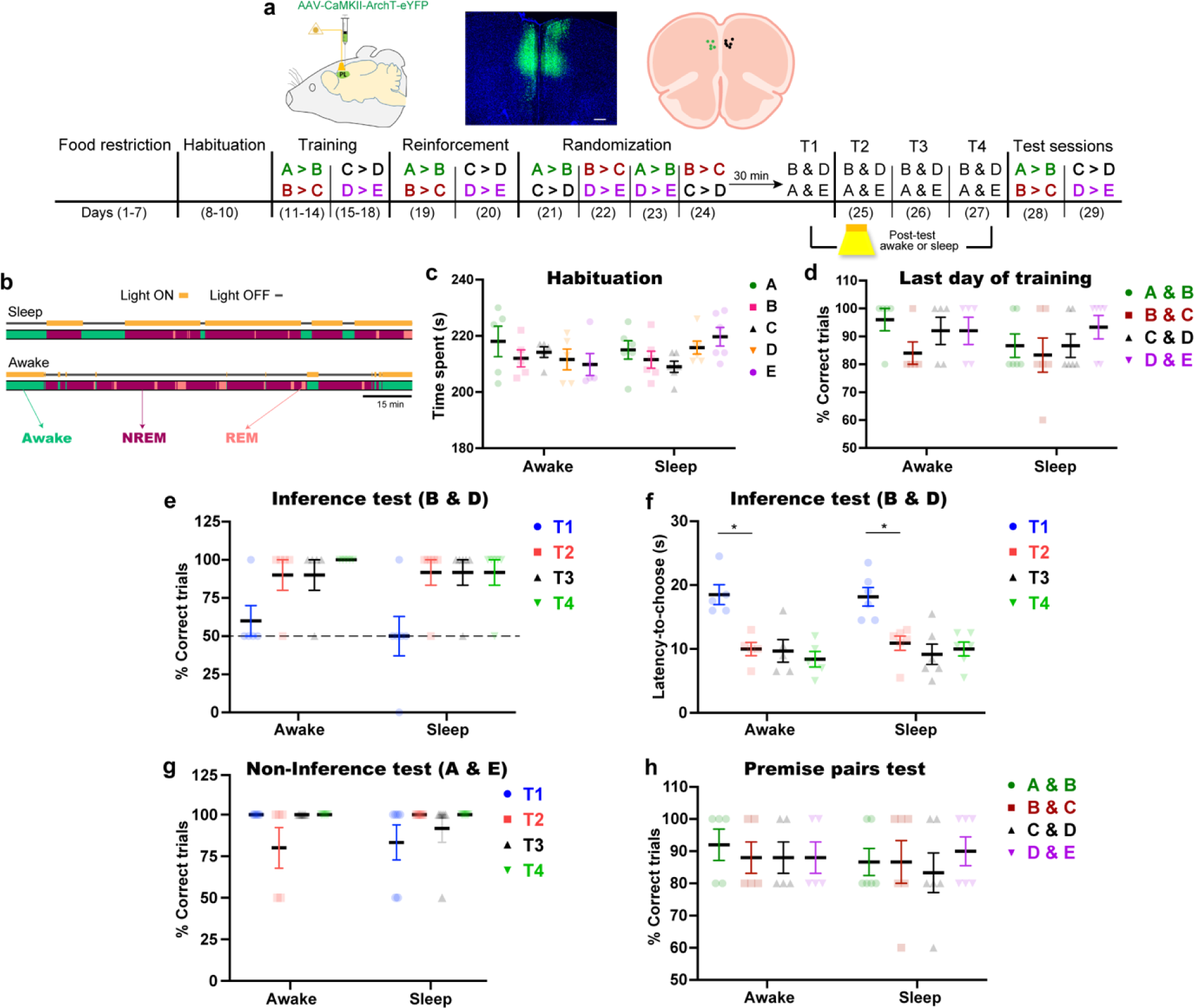
Prelimbic cortex activity is not required for the emergence of inference. **a,** Top, Labelling of excitatory neurons of the prelimbic cortex (PL) with ArchT (left), coronal section from the mouse brain showing the expression of ArchT-eYFP (green) in the PL (middle) and optic fibre tip location (right). Black dots, mice in the awake group; Green dots, mice in the sleep group. Blue, 4′,6-diamidino-2-phenylindole (DAPI) staining. Scale bar, 100 µm. Bottom, The behavioural schedule used to manipulate PL activity during sleep or awake periods after test sessions. The order of presentation of the premise pairs during the randomization stage is different across animals within the same group (See Methods; Table 1). **b**, Diagram showing the state-specific manipulation. Scale bar, 15 minutes. **c,** Time spent in each context during the habituation phase**. d,** Performance during the last day of training (day 14 for A > B and B > C; day 18 for C > D and D > E) for each premise pair. **e-h,** Performance during the test sessions; the percent of correct trials during the inference test (**e**), latency time to choose during the inference test (**f**), during the non-inference test (**g**), during the original premise pairs test (**h)**. T1, test 1; T2, test 2; T3, test 3; T4, test 4. *n* = 5 mice for the awake group; *n* = 6 mice for the sleep group. Statistical comparisons were made using a two-way repeated-measures analysis of variance (ANOVA) with Tukey’s multiple comparison test (**c-h**). In the non-inference test (**g**), the statistical significance denotes the comparison between performance of the two groups relative to the chance level (50%). \**P* < 0.05; \*\*\*\**P* < 0.0001; ns, not significant (*P* > 0.05). Data are presented as the mean ± standard error of the mean (s.e.m.).

**Extended Data Fig. 6.**
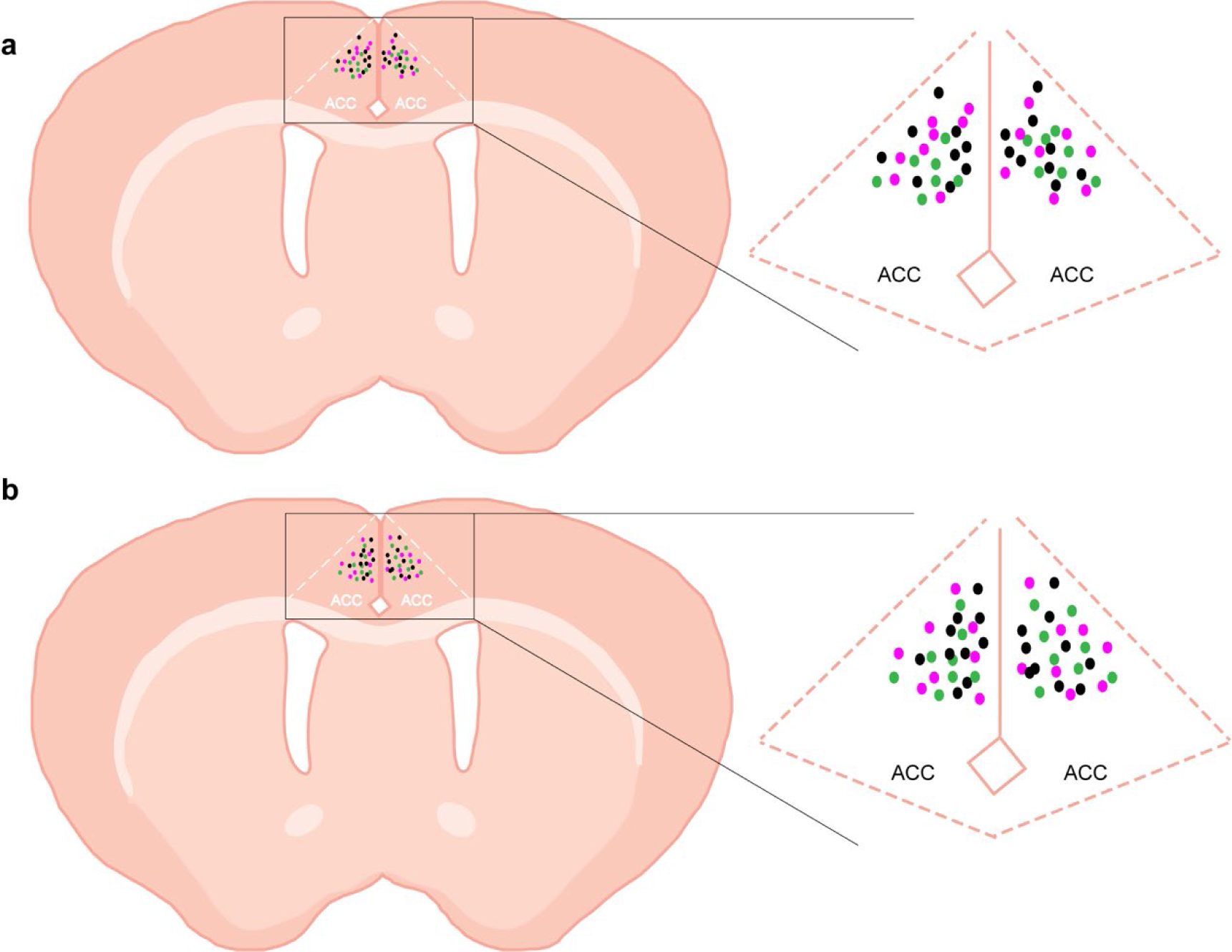
Optic fibre locations in mice used in the manipulation experiments. **a,** Coronal section from the mouse brain showing the optic fibre tip location for mice shown in Fig. 2. Black dots, mice in the awake group; Green dots, mice in the NREM group; Pink dots, mice in the REM group. **b**, Coronal section from the mouse brain showing the optic fibre tip location for mice shown in Fig. 5. Black dots, mice in the light off group; Green dots, mice in the NREM group; Pink dots, mice in the REM group. The anterior cingulate cortex (ACC) is indicated by dashed lines.

**Extended Data Fig. 7.**
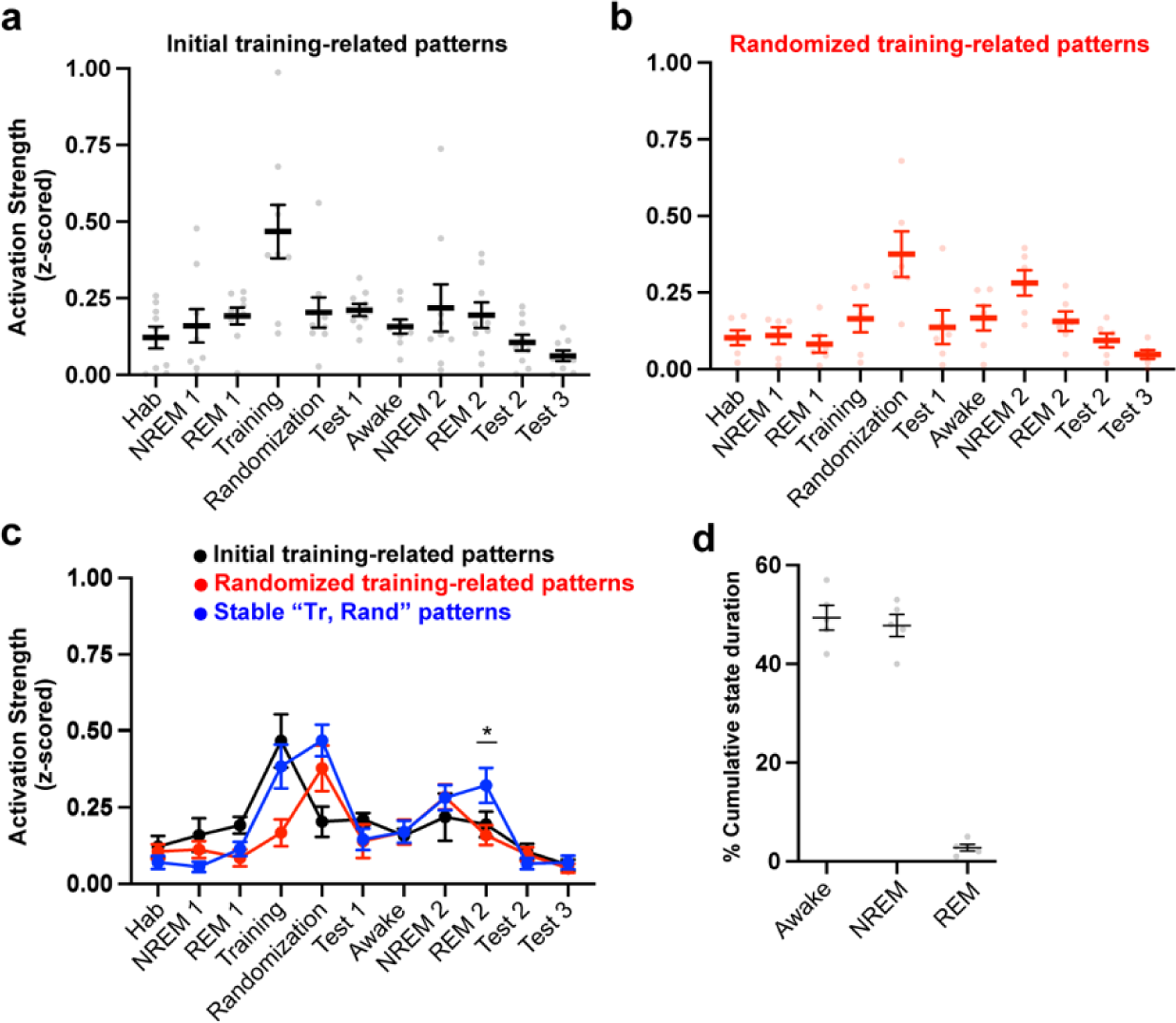
Temporally different patterns representing the original premise pairs. **a-c**, Activation strength (z-scored) of the patterns representing the original premise pairs during initial training phase (**a, c**), during randomized training phase (**b, c**). **d,** Cumulative duration of each state (awake and sleep states) in all 5 mice. Statistical comparison was done using unpaired t-test; *P < 0.05. Data are presented as the mean ± standard error of the mean (s.e.m.).

**Extended Data Fig. 8.**
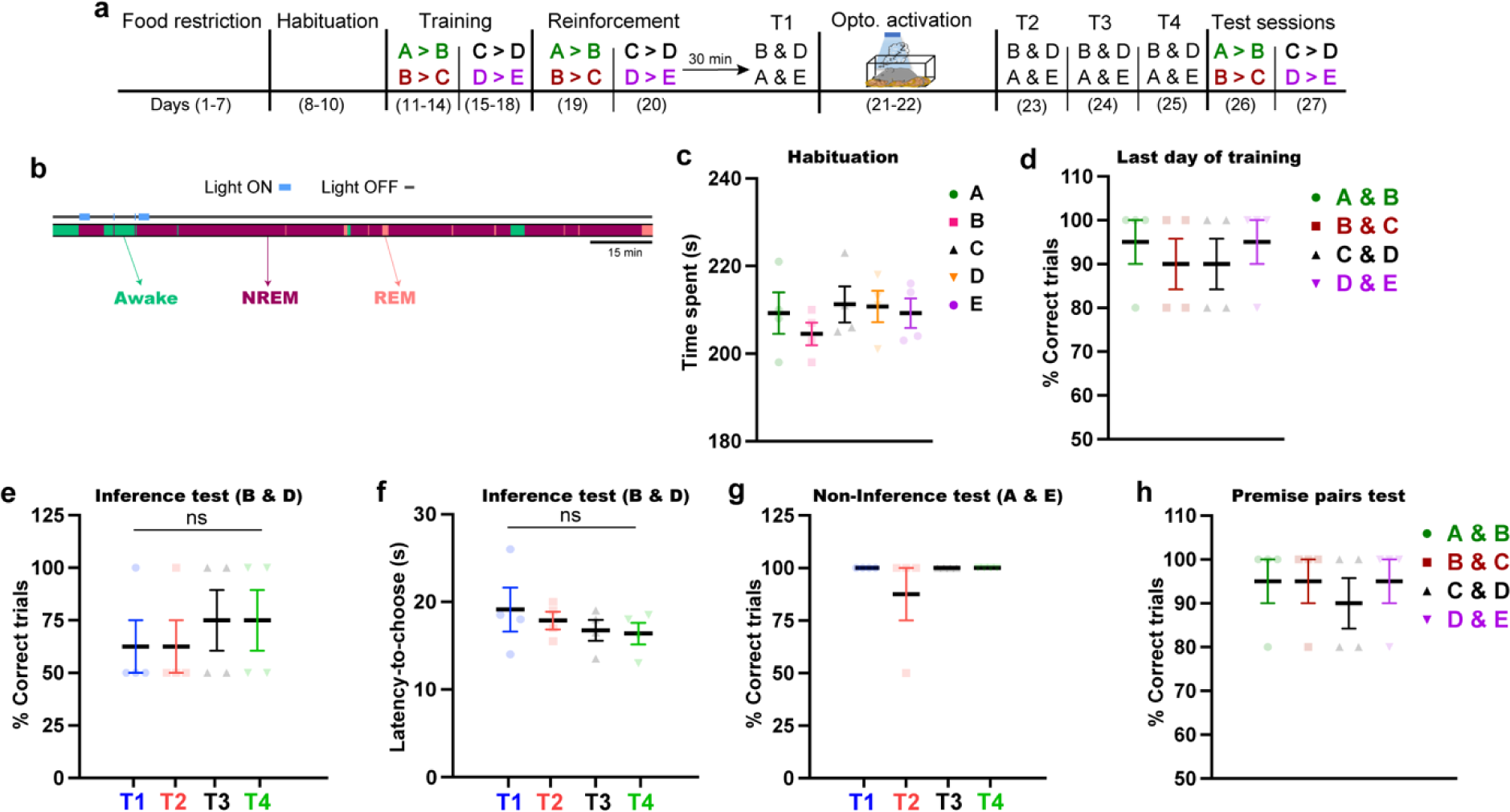
Optogenetic activation of MEC→ACC circuit during NREM sleep is not sufficient for the emergence of inference. **a,** The behavioural schedule used to manipulate MEC→ACC circuit activity during NREM sleep after test sessions. **b**, Diagram showing the NREM-specific manipulation. The manipulation protocol mimicked the protocol used during REM sleep manipulation in Fig. 5 (see Methods) to avoid the prolonged light stimulation which might affect the results. Scale bar, 15 minutes. **c**, The time spent in each context during the habituation phase. **d**, Performance during the last day of training (day 14 for A > B and B > C; day 18 for C > D and D > E) for each premise pair. **e**-**h**, Performance during the test sessions; the percent of correct trials during the inference test (**e**), latency time to choose during the inference test (**f**), during the non-inference test (**g**), during the original premise pairs test (**h**). T1, test 1; T2, test 2; T3, test 3; T4, test 4; *n* = 4 mice. Statistical comparisons were made using a one-way repeated-measures analysis of variance (ANOVA) with Tukey’s multiple comparison test (**c**-**h**). ns, not significant (*P* > 0.05). Data are presented as the mean ± standard error of the mean (s.e.m.).

**Extended Data Fig. 9.**
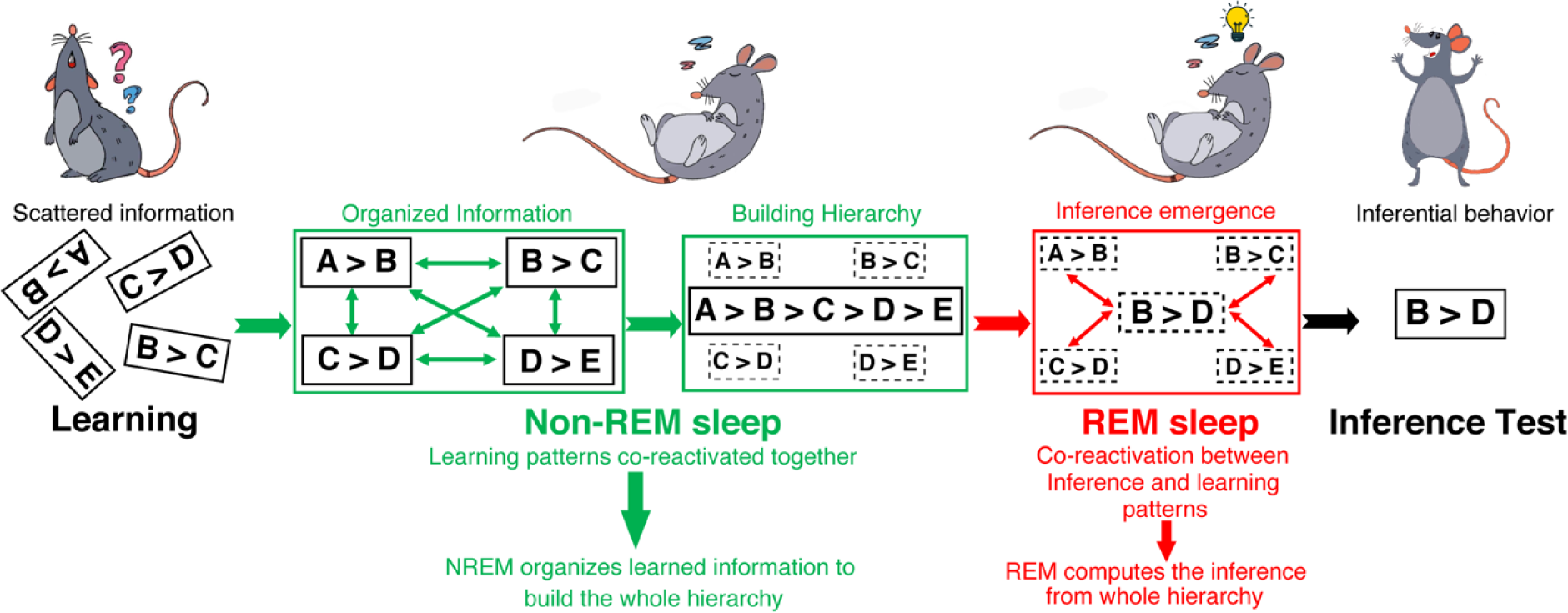
A model of the coordinated roles of NREM & REM sleep in building inferential knowledge. Summary of the process of inference emergence. It starts with learning temporally distinct, but related information. Then, systematic organization of learned information in hierarchy during NREM sleep. Afterwards, building the inferential information from the hierarchy during REM sleep. Subsequently, inferential behaviour appears in the test session. An arrow with arrowheads at both ends indicates synchronous activity.

## References

1 Buzsaki, G. & Fernandez-Ruiz, A. Utility of the Idling Brain: Abstraction of New Knowledge. Cell 178, 513–515 (2019). 10.1016/j.cell.2019.07.004

2 Ghandour, K. & Inokuchi, K. Memory reactivations during sleep. Neurosci Res 189, 60–65 (2023). 10.1016/j.neures.2022.12.018

3 Ellenbogen, J. M., Hu, P. T., Payne, J. D., Titone, D. & Walker, M. P. Human relational memory requires time and sleep. Proc Natl Acad Sci U S A 104, 7723–7728 (2007). 10.1073/pnas.0700094104

4 Wagner, U., Gais, S., Haider, H., Verleger, R. & Born, J. Sleep inspires insight. Nature 427, 352–355 (2004). 10.1038/nature02223

5 Behrens, T. E. J. et al. What Is a Cognitive Map? Organizing Knowledge for Flexible Behavior. Neuron 100, 490–509 (2018). 10.1016/j.neuron.2018.10.002

6 Brunamonti, E. et al. Neuronal Modulation in the Prefrontal Cortex in a Transitive Inference Task: Evidence of Neuronal Correlates of Mental Schema Management. The Journal of neuroscience : the official journal of the Society for Neuroscience 36, 1223–1236 (2016). 10.1523/JNEUROSCI.1473-15.2016

7 Spalding, K. N. et al. Ventromedial Prefrontal Cortex Is Necessary for Normal Associative Inference and Memory Integration. The Journal of neuroscience : the official journal of the Society for Neuroscience 38, 3767–3775 (2018). 10.1523/JNEUROSCI.2501-17.2018

8 Ghandour, K. et al. Orchestrated ensemble activities constitute a hippocampal memory engram. Nat Commun 10, 2637 (2019). 10.1038/s41467-019-10683-2

9 Wilson, M. A. & McNaughton, B. L. Reactivation of hippocampal ensemble memories during sleep. Science 265, 676–679 (1994). 10.1126/science.8036517

10 Boyce, R., Glasgow, S. D., Williams, S. & Adamantidis, A. Causal evidence for the role of REM sleep theta rhythm in contextual memory consolidation. *Science (New York*, N.Y.) 352, 812–816 (2016). 10.1126/science.aad5252

11 Girardeau, G., Benchenane, K., Wiener, S. I., Buzsaki, G. & Zugaro, M. B. Selective suppression of hippocampal ripples impairs spatial memory. Nat Neurosci 12, 1222–1223 (2009). 10.1038/nn.2384

12 Gridchyn, I., Schoenenberger, P., O’Neill, J. & Csicsvari, J. Assembly-Specific Disruption of Hippocampal Replay Leads to Selective Memory Deficit. Neuron 106, 291–300 e296 (2020). 10.1016/j.neuron.2020.01.021

13 Diekelmann, S. & Born, J. The memory function of sleep. Nat Rev Neurosci 11, 114–126 (2010). 10.1038/nrn2762

14 Gupta, A. S., van der Meer, M. A., Touretzky, D. S. & Redish, A. D. Hippocampal replay is not a simple function of experience. Neuron 65, 695–705 (2010). 10.1016/j.neuron.2010.01.034

15 Olafsdottir, H. F., Barry, C., Saleem, A. B., Hassabis, D. & Spiers, H. J. Hippocampal place cells construct reward related sequences through unexplored space. eLife 4, e06063 (2015). 10.7554/eLife.06063

16 Olafsdottir, H. F., Bush, D. & Barry, C. The Role of Hippocampal Replay in Memory and Planning. Current biology : CB 28, R37–R50 (2018). 10.1016/j.cub.2017.10.073

17 Cousins, J. N., El-Deredy, W., Parkes, L. M., Hennies, N. & Lewis, P. A. Cued memory reactivation during slow-wave sleep promotes explicit knowledge of a motor sequence. The Journal of neuroscience : the official journal of the Society for Neuroscience 34, 15870–15876 (2014). 10.1523/JNEUROSCI.1011-14.2014

18 Gais, S. & Born, J. Low acetylcholine during slow-wave sleep is critical for declarative memory consolidation. Proc Natl Acad Sci U S A 101, 2140–2144 (2004). 10.1073/pnas.0305404101

19 Plihal, W. & Born, J. Effects of early and late nocturnal sleep on declarative and procedural memory. J Cogn Neurosci 9, 534–547 (1997). 10.1162/jocn.1997.9.4.534

20 Plihal, W. & Born, J. Memory consolidation in human sleep depends on inhibition of glucocorticoid release. Neuroreport 10, 2741–2747 (1999). 10.1097/00001756-199909090-00009

21 Walker, M. P. The role of sleep in cognition and emotion. Annals of the New York Academy of Sciences 1156, 168–197 (2009). 10.1111/j.1749-6632.2009.04416.x

22 Wamsley, E. J., Tucker, M., Payne, J. D., Benavides, J. A. & Stickgold, R. Dreaming of a learning task is associated with enhanced sleep-dependent memory consolidation. Current biology : CB 20, 850–855 (2010). 10.1016/j.cub.2010.03.027

23 Cai, D. J., Mednick, S. A., Harrison, E. M., Kanady, J. C. & Mednick, S. C. REM, not incubation, improves creativity by priming associative networks. Proceedings of the National Academy of Sciences of the United States of America 106, 10130–10134 (2009). 10.1073/pnas.0900271106

24 Poe, G. R., Walsh, C. M. & Bjorness, T. E. Cognitive neuroscience of sleep. Prog Brain Res 185, 1–19 (2010). 10.1016/B978-0-444-53702-7.00001-4

25 Brodt, S., Inostroza, M., Niethard, N. & Born, J. Sleep-A brain-state serving systems memory consolidation. Neuron 111, 1050–1075 (2023). 10.1016/j.neuron.2023.03.005

26 Lewis, P. A., Knoblich, G. & Poe, G. How Memory Replay in Sleep Boosts Creative Problem-Solving. Trends in cognitive sciences 22, 491–503 (2018). 10.1016/j.tics.2018.03.009

27 Bramham, C. R. & Srebro, B. Synaptic plasticity in the hippocampus is modulated by behavioral state. Brain Res 493, 74–86 (1989). 10.1016/0006-8993(89)91001-9

28 Basheer, R. et al. Effects of sleep on wake-induced c-fos expression. J Neurosci 17, 9746–9750 (1997).

29 Cirelli, C. & Tononi, G. Gene expression in the brain across the sleep-waking cycle. Brain Res 885, 303–321 (2000). 10.1016/s0006-8993(00)03008-0

30 Cirelli, C. & Tononi, G. Differential expression of plasticity-related genes in waking and sleep and their regulation by the noradrenergic system. J Neurosci 20, 9187–9194 (2000).

31 Pompeiano, M., Cirelli, C., Arrighi, P. & Tononi, G. c-Fos expression during wakefulness and sleep. Neurophysiol Clin 25, 329–341 (1995). 10.1016/0987-7053(96)84906-9

32 Pompeiano, M., Cirelli, C., Ronca-Testoni, S. & Tononi, G. NGFI-A expression in the rat brain after sleep deprivation. Brain Res Mol Brain Res 46, 143–153 (1997). 10.1016/s0169-328x(96)00295-1

33 Pompeiano, M., Cirelli, C. & Tononi, G. Immediate-early genes in spontaneous wakefulness and sleep: expression of c-fos and NGFI-A mRNA and protein. J Sleep Res 3, 80–96 (1994). 10.1111/j.1365-2869.1994.tb00111.x

34 Ackermann, S. & Rasch, B. Differential effects of non-REM and REM sleep on memory consolidation? Curr Neurol Neurosci Rep 14, 430 (2014). 10.1007/s11910-013-0430-8

35 Tamaki, M. et al. Complementary contributions of non-REM and REM sleep to visual learning. Nat Neurosci 23, 1150–1156 (2020). 10.1038/s41593-020-0666-y

36 Pereira, S. I. R. & Lewis, P. A. The differing roles of NREM and REM sleep in the slow enhancement of skills and schemas. Current Opinion in Physiology 15, 82–88 (2020). 10.1016/j.cophys.2019.12.005

37 Preston, A. R. & Eichenbaum, H. Interplay of hippocampus and prefrontal cortex in memory. Current biology : CB 23, R764–773 (2013). 10.1016/j.cub.2013.05.041

38 Barron, H. C. et al. Neuronal Computation Underlying Inferential Reasoning in Humans and Mice. Cell 183, 228–243 e221 (2020). 10.1016/j.cell.2020.08.035

39 Zeithamova, D., Dominick, A. L. & Preston, A. R. Hippocampal and ventral medial prefrontal activation during retrieval-mediated learning supports novel inference. Neuron 75, 168–179 (2012). 10.1016/j.neuron.2012.05.010

40 Miyamoto, D. et al. Top-down cortical input during NREM sleep consolidates perceptual memory. Science 352, 1315–1318 (2016). 10.1126/science.aaf0902

41 Takekawa, T. et al. Automatic sorting system for large calcium imaging data. bioRxiv, 215145 (2017). 10.1101/215145

42 Aly, M. H., Abdou, K., Okubo-Suzuki, R., Nomoto, M. & Inokuchi, K. Selective engram coreactivation in idling brain inspires implicit learning. Proc Natl Acad Sci U S A 119, e2201578119 (2022). 10.1073/pnas.2201578119

43 Dusek, J. A. & Eichenbaum, H. The hippocampus and memory for orderly stimulus relations. Proc Natl Acad Sci U S A 94, 7109–7114 (1997). 10.1073/pnas.94.13.7109

44 Buckmaster, C. A., Eichenbaum, H., Amaral, D. G., Suzuki, W. A. & Rapp, P. R. Entorhinal cortex lesions disrupt the relational organization of memory in monkeys. J Neurosci 24, 9811–9825 (2004). 10.1523/JNEUROSCI.1532-04.2004

45 Liu, Y., Dolan, R. J., Kurth-Nelson, Z. & Behrens, T. E. J. Human Replay Spontaneously Reorganizes Experience. Cell 178, 640–652 e614 (2019). 10.1016/j.cell.2019.06.012

46 Hasegawa, E. et al. Rapid eye movement sleep is initiated by basolateral amygdala dopamine signaling in mice. Science 375, 994–1000 (2022). 10.1126/science.abl6618

47 Fujisawa, S. & Buzsaki, G. A 4 Hz oscillation adaptively synchronizes prefrontal, VTA, and hippocampal activities. Neuron 72, 153–165 (2011). 10.1016/j.neuron.2011.08.018

48 Walker, M. P., Liston, C., Hobson, J. A. & Stickgold, R. Cognitive flexibility across the sleep-wake cycle: REM-sleep enhancement of anagram problem solving. Brain Res Cogn Brain Res 14, 317–324 (2002). 10.1016/s0926-6410(02)00134-9

49 Konno, A. & Hirai, H. Efficient whole brain transduction by systemic infusion of minimally purified AAV-PHP.eB. J Neurosci Methods 346, 108914 (2020). 10.1016/j.jneumeth.2020.108914

50 Dana, H. et al. High-performance calcium sensors for imaging activity in neuronal populations and microcompartments. Nat Methods 16, 649–657 (2019). 10.1038/s41592-019-0435-6

51 Iida, A., Takino, N., Miyauchi, H., Shimazaki, K. & Muramatsu, S. Systemic delivery of tyrosine-mutant AAV vectors results in robust transduction of neurons in adult mice. Biomed Res Int 2013, 974819 (2013). 10.1155/2013/974819

52 Li, X. G. et al. Viral-mediated temporally controlled dopamine production in a rat model of Parkinson disease. Mol Ther 13, 160–166 (2006). 10.1016/j.ymthe.2005.08.009

53 Nomoto, M. et al. Hippocampus as a sorter and reverberatory integrator of sensory inputs. Nat Commun 13, 7413 (2022). 10.1038/s41467-022-35119-2

54 Middleton, S. J. et al. Altered hippocampal replay is associated with memory impairment in mice heterozygous for the Scn2a gene. Nat Neurosci 21, 996–1003 (2018). 10.1038/s41593-018-0163-8

55 Trouche, S. et al. Recoding a cocaine-place memory engram to a neutral engram in the hippocampus. Nat Neurosci 19, 564–567 (2016). 10.1038/nn.4250

56 Watrous, A. J., Miller, J., Qasim, S. E., Fried, I. & Jacobs, J. Phase-tuned neuronal firing encodes human contextual representations for navigational goals. Elife 7 (2018). 10.7554/eLife.32554

